# Dynamics of sphingolipids and the serine-palmitoyltransferase complex in oligodendrocytes during myelination

**DOI:** 10.1101/2020.01.15.908277

**Authors:** Deanna L. Davis, Usha Mahawar, Victoria S. Pope, Jeremy Allegood, Carmen Sato-Bigbee, Binks W. Wattenberg

## Abstract

Myelin is a unique, lipid-rich membrane structure that accelerates neurotransmission and supports neuronal function. Sphingolipids are critical components of myelin. Here we examined sphingolipid synthesis during the peak period of myelination in the postnatal rat brain. Importantly, we made measurements in isolated oligodendrocytes, the myelin-producing cells in the central nervous system. We analyzed sphingolipid distribution and levels of critical enzymes and regulators in the sphingolipid biosynthetic pathway, with a focus on the serine palmitoyltransferase (SPT) complex, the rate-limiting step in this pathway. During myelination levels of the major SPT subunits increased and oligodendrocyte maturation was accompanied by extensive alterations in the composition of the SPT complex. These included changes in the relative levels of alternate catalytic subunits, SPTLC2 and −3, the relative levels of isoforms of the small subunits ssSPTa and –b, and in the isoform distribution of the SPT regulators, the ORMDLs. As myelination progressed there were distinct changes in both the nature of the sphingoid backbone and the N-acyl chains incorporated into sphingolipids. The distribution of these changes among sphingolipid family members indicates that there is selective channeling of the ceramide backbone towards specific downstream metabolic pathways during myelination.

## Introduction

Myelin is a highly specialized membrane that wraps around axons in the central and peripheral nervous systems, accelerating neuronal transmission and sustaining neuronal function and viability. Loss of myelination is the pathological basis for several devastating neurological disorders, collectively known as demyelinating diseases, the most common of which is multiple sclerosis (1). The myelin membrane is an extension of the plasma membrane of specialized cells; oligodendrocytes in the central nervous system and Schwann cells in the peripheral nervous system. The composition of myelin is highly distinct from other biological membranes both with regard to protein and lipid composition. Lipids compose approximately 80% (by weight) of myelin, as compared to 50% or less in most other biological membranes (2). The lipid portion of myelin includes glycerolipids, sterol, and remarkably high levels of sphingolipids. One recent report determined that 80%of the non-sterol lipid in myelin was sphingolipid (3). The majority of myelin sphingolipid is cerebroside, the term for the monoglycosylated ceramides glucosylceramide and galactosylceramide, and sulfatide, which is a sulfated galactosyl ceramide. Myelin also contains substantial amounts of sphingomyelin. Myelin sphingolipids determine the unique physical characteristics of this membrane (reviewed in (4)) and promote membrane interactions within the myelin structure and between myelin and neurons (5-9). The importance of myelin sphingolipids is confirmed by the observation that mice deficient in the production of galactosylceramide have profound defects in myelination (10,11) and deletion of sulfatide yields defects in myelin structure that become more severe as mice mature (8,12).

The production of all sphingolipids begins with the initiating and rate limiting enzyme in the pathway, serine palmitoyltransferase (SPT), a protein complex embedded in the endoplasmic reticulum (reviewed in (13)). Most commonly this enzyme condenses serine with palmitoyl-coenzyme A (CoA) to produce 3-ketodihydrosphingosine, an 18 carbon precursor of all sphingolipids. It is expected that activity of this enzyme would be a key regulatory step for sphingolipid biosynthesis for myelin production. SPT is a multi-subunit enzyme which has a core composed of two subunits, most commonly SPTLC1 and −2. SPTLC2 contains the active site of the enzyme, but is not active by itself, requiring assembly with SPTLC1 for activity and stability. In the core dimer SPTLC2 can be replaced by a homologous subunit, SPTLC3. SPT activity is strongly affected by accessory subunits. The so-called small SPT subunits, ssSPTa and –b, strongly enhance activity of the core SPT complex. In addition to determining the overall level of SPT activity, the subunit composition of SPT specifies which fatty-acyl-CoA substrates can be utilized (14). The complex formed by SPTLC1/2/ssSPTa utilizes the 16 carbon acyl-CoA palmitoyl-CoA and therefore produces 18 carbon sphingolipids, which are the most abundant species. But SPTLC3 and/or ssSPTb, depending on the combination, can allow the use of both shorter and longer acyl-CoAs and therefore produce less common sphingoid bases. The functional effects of these alterations are not yet clear, but changes in SPT complex composition, as we find here, suggests that these changes may fine-tune the characteristics of the sphingolipid component of myelin. SPT is homeostatically regulated; SPT activity is diminished when cellular sphingolipid levels rise (15,16). This control requires the ORMDLs, membrane-bound proteins of the endoplasmic reticulum which form stable complexes with SPT (Reviewed in (17)). Mammalian cells contain three ORMDL isoforms whose individual functions have yet to be well characterized. The ORMDL/SPT complex responds to ceramide levels. One major function of this regulation may be to control sphingolipid biosynthesis to maintain the ceramide pool at sub-apoptotic levels.

Once SPT produces 3-keto dihyrosphingosine, this molecule is reduced to dihydrosphingosine which may then be acylated by one of the six ceramide synthases to produce dihydroceramide, which is then desaturated by ceramide desaturase to produced ceramide, the central substrate for higher order sphingolipids (reviewed in (18)). Each ceramide synthase has a distinct preference for acyl-CoA substrates of different chain lengths, typically from 14 to 26 carbons in length, and therefore a variety of ceramide molecular species are produced. Ceramide is the backbone for sphingomyelin, by the transfer of phosphocholine from phosphatidylcholine by the sphingomyelin synthases. Important for myelin production, ceramide is also the backbone of glycosphingolipids (reviewed in (19)). The synthesis of glycosphingolipids is initiated by the transfer of either glucose or galactose to ceramide by UDP-glucose ceramide glucosyltransferase (UGCG) or UDP-glycosyltransferase 8 (UGT8) respectively. Galactosylceramide is sulfated by galactose-3-O-sulfotransferase (Gal3st) to produce sulfatide. Ceramide can also be deacylated by one of several ceramidases to generate sphingosine (reviewed in (20)). Sphingosine and dihydrosphingosine may be phosphorylated by one of the two sphingosine kinases to produce the potent signaling molecules sphingosine- or dihydrosphingosine-1-phosphate, both of which are substrates for the only enzyme that can degrade the sphingosine backbone, sphingosine-1-phosphate lyase (reviewed in (21)).

Here we analyze the well characterized rat brain system during the post-natal peak of myelination with respect to levels of sphingolipids and sphingolipid metabolic enzymes. Importantly, for the first time we analyze these components in isolated oligodendrocytes to gain direct insight into myelin production that is not possible when analyzing total brain samples. Our aim is to establish the basis for elucidating the regulatory mechanisms that determine both the timing and lipid specificity of myelin sphingolipid production under normal physiological conditions and how those mechanisms may be disturbed in demyelinating diseases.

## 2. Materials and Methods

### 2.1 Materials

The following primary antibodies were used in this study: anti-MBP (#MAB386), anti-MOG (#MAB5680), anti-MAG (#MAB1567), and anti-SPTLC2 (#ABS1641) were from EMD Millipore (Temecula, CA); anti-ORMDL (#NBP1-98511) and anti-GAPDH (#NB300-221) were from Novus (Centennial, CO); anti-SPTLC1 (#611305) was from BD Transduction Labs (San Jose, CA); anti-SPTLC3 (#PA5-65493) was from Invitrogen/ThermoFisher (Waltham, MA); anti-PLP (#ab28486) was from Abcam (Cambridge, MA); anti-Calnexin (#ADI-SPA-860) was from Enzo (Farmingdale, NY). Amersham ECL Prime Western Blot Detection Reagent (#RPN2322) was from GE Healthcare (Marlborough, MA). PVDF was from BioRad (Hercules, CA). HRP-conjugated secondary antibodies for mouse (#31430) and rabbit (#31460) were purchased from Thermo-Fisher (Waltham, MA) and goat anti-rat (#112-035-003) was from Jackson Immuno Research Labs (West Grove, PA).

Quantitative real-time PCR reagents used in this study were from IDT (San Jose, CA) as follows: Prime Time Gene Expression Master Mix (#1055772) and FAM-labeled PrimeTime qPCR 5’ nuclease probes for rat genome (gene ID# and sequence information for each primer/probe set is included in *Supplementary Information*). Thin-wall PCR plates (#HSP9601) and Microseal optical plate covers (#MSB1001) were from BioRad (Hercules, CA). Trizol Reagent (#15596026) was from Invitrogen/Thermo (Waltham, MA). 5 Prime heavy 2 mL-phase lock gels (#2302830) were from VWR (Radnor, PA). High Capacity cDNA Reverse Transcription Kit (#4368814) was from ABI/Thermo (Waltham, MA).

Internal standards for mass spectrometry were purchased from Avanti Polar Lipids (Alabaster, AL). Internal standards were added to samples in 10 uL ethanol:methanol:water (7:2:1) as a cocktail of 250 pmol each. Standards for sphingoid bases and sphingoid base 1-phosphates were 17-carbon chain length analogs: C17-sphingosine, (2S,3R,4E)-2-aminoheptadec-4-ene-1,3-diol (d17:1-So); C17-sphinganine, (2S,3R)-2-aminoheptadecane-1,3-diol (d17:0-Sa); C17-sphingosine 1-phosphate, heptadecasphing-4-enine-1-phosphate (d17:1-So1P); and C17-sphinganine 1-phosphate, heptadecasphinganine-1-phosphate (d17:0-Sa1P). Standards for N-acyl sphingolipids were C12-fatty acid analogs: C12-Cer, N-(dodecanoyl)-sphing-4-enine (d18:1/C12:0); C12-Cer 1-phosphate, N-(dodecanoyl)-sphing-4-enine-1-phosphate (d18:1/C12:0-Cer1P); C12-sphingomyelin, N-(dodecanoyl)-sphing-4-enine-1-phosphocholine (d18:1/C12:0-SM); and C12-glucosylceramide, N-(dodecanoyl)-1-β-glucosyl-sphing-4-eine

Deuterated L-serine (#DLM-161-PK) was from Cambridge Isotope Labs (Tewksbury, MA) and myriocin (#63150) was from Cayman Chemicals (Ann Arbor, MI). Percoll (#17-0891-01) was from GE Healthcare (Marlborough, MA). HBSS (#14185-052) and DMEM/F12 (#11330-032) were from Gibco/Thermo (Waltham, MA). Bovine pancreas DNAse (DN-25) and papain (#P-4762) were from Sigma-Aldrich (St. Louis, MO). Protease inhibitor cocktail (Complete, EDTA-free) was from Roche (Indianapolis, IN). Unless otherwise indicated all remaining reagents used were from commercial sources and readily available.

### 2.2 Animals

Sprague-Dawley female rats and their natural 2-day-old pups (10 pups/litter) were obtained from Charles River Laboratories (Wilmington, MA). Animals were housed under light/dark cycle and temperature controlled conditions and allowed food and water *ad libitum*. Studies were conducted in accordance with the National Institutes of Health Guidelines for the Care and Use of Animals in Research and under protocols approved by the Animal Care and Use Committee of Virginia Commonwealth University (VCU), Richmond, VA.

### 2.3 Isolation of primary oligodendrocytes from rat brain

Oligodendrocytes were freshly isolated from rat brain at varying times after birth (Day 2, Day 8 and Day 16) using a Percoll gradient centrifugation and differential cell attachment protocol (adapted from (22)). Cells isolated at postnatal day 2 are proliferating immature oligodendrocyte progenitors while those from 8 day-old pups are postmitotic pre-oligodendrocytes, a developmental stage that immediately precedes their differentiation into mature myelinating cells. Cells isolated from 16-day-old animals are in their great majority fully differentiated oligodendrocytes. Animals were rapidly decapitated per guidelines set forth by IACUC of VCU. The meninges were removed from the dissected cerebral hemispheres and brain tissue was placed in ice-cold Buffer A (25mM HEPES in HBSS, 1 mg/mL Glucose, pH 7.2). After all tissue was collected, the brains were finely minced then placed in Buffer A containing 10μg/mL bovine pancreas DNAse (Sigma #DN25) and 1Unit/mL papain (Sigma #P4762). Tissue was transferred to a sterile beaker then digested for 25 minutes in a 37°C shaking water bath set at 280rpm. Following digestion, tissue was transferred to 50mL-conical tubes and enzyme activity was quenched by dilution with additional ice-cold buffer A containing DNAse. Samples were centrifuged at 500 rpm for 5 minutes at 4°C then washed three times with maximal volumes of cold buffer A. The washed, digested tissue was passed through a sterile nylon mesh (75μm, SEFAR, Buffalo, NY) to create a single cell suspension. Filtrates were diluted with buffer A then added to 15ml Corex glass tubes containing isotonic Percoll at a ratio of 1.5:1 (cells:Percoll, v/v). Tubes were capped then centrifuged for 15 minutes at 30,000x*g* in a JLA 16.250 fixed angle rotor (Beckman-Coulter, Brea, CA) at 4°C using a Beckman Avanti JE centrifuge. Oligodendrocytes were carefully isolated from the Percoll gradient, pelleted, washed once with Buffer A to remove residual Percoll, then resuspended in DMEM/Ham’s F12 media containing 1mg/mL fatty acid free BSA. The cells were placed in a 100mm tissue culture dish treated Petri dish and incubated at 37°C for 20 minutes to allow for differential attachment of microglia and potential remaining astrocytes. After gently swirling, the floating oligodendrocytes were pelleted and then divided into equal aliquots for RNA and protein isolation protocols.

### 2.4 Preparation of total membranes from whole rat cerebral hemispheres

Membrane isolation used a procedure adapted from (23). For each postnatal age group reported in this study, brains were harvested from five rat pups and stored in separate tubes. Immediately upon harvesting, the cerebral hemisphere brain tissue was snap frozen using liquid nitrogen and stored at −80°C until time of processing. Frozen cerebral hemispheres were placed into 5 mL ice-cold Buffer A (20mM HEPES-KOH (pH 7.0), 320mM Sucrose, 1X protease inhibitor cocktail) in a 7-mL glass dounce tissue homogenizer (Kontes, Kimble/Chase, Rockwood, TN) and then homogenized using the ‘A’ pestle (4 strokes) followed by the ‘B’ pestle (8 strokes). Samples were transferred to 15-mL centrifuge tubes on ice then centrifuged at 1000*xg* for 10 minutes at 4°C to clear nuclei and cell debris. Supernatants were transferred to Opti-Seal tubes (Beckman #361621), placed in an SLA110 rotor and centrifuged at 100K rpm (Beckman-Coulter Optima Max XP) for 30 minutes at 4°C to collect total membranes. Membrane pellets were resuspended in Buffer A and then passaged approximately 10X through a 26 gauge needle. Total protein was quantitated using the Bradford protein assay (Biorad, Hercules, CA) and aliquots were stored at −80°C until processing for Western blot, SPT activity assays or lipidomics analysis.

### 2.5 Isolation of total RNA from whole rat cerebral hemispheres

Immediately upon harvesting, half of the cerebral hemisphere brain tissue from each rat pup was placed into Trizol Reagent and then snap frozen using liquid nitrogen and stored at −80°C until time of processing. Frozen Trizol/brain tissue samples were thawed on ice, then transferred to a 7-mL glass dounce and homogenized until smooth. This mixture was transferred to a pre-spun 2-mL heavy phase lock gel (5 Prime) after which 200μL of chloroform/isoamyl alcohol (49:1, v/v, Sigma cat# 25668) was added and samples were vortexed vigorously. After standing at room temperature for approximately 3 minutes, phases were separated by centrifuging at 9000 rpm for 15 minutes at 4°C. The upper (aqueous) phase was transferred to a sterile 1.5 mL microfuge tube and 500μL isopropanol was added to each tube to precipitate RNA. RNA was pelleted by centrifugation at full speed for 20 minutes using a refrigerated table-top centrifuge. RNA pellets were washed three times with 70% ethanol, then dried under sterile conditions. Pellets were resuspended using RNAse/DNAse-free water (50-100μL depending on pellet size). RNA content was quantitated using a NanoDrop 2000, and aliquots were prepared and stored at −80°C until further use. To assess the quality of the isolated RNA prior to use, 1μg of each sample was resolved using 1% agarose gels containing 1% Clorox™ bleach (24)(*Electrophoresis*, 2012; 33(2): 366–369).

### 2.6 cDNA synthesis and quantitative PCR analysis

Single-stranded cDNA was synthesized using 2μg of total RNA isolated from the brain tissue of individual rat pups. We used the High Capacity cDNA Reverse Transcription Kit (#4368814) from ABI/Thermo Fisher per the manufacturer’s instructions. FAM-labeled qPCR primer-probe sets were designed for each gene of interest and ordered from IDT (see *Supplementary Information*). Real-time qPCR analysis was performed using 30-50ng of cDNA per animal per gene combined with PrimeTime Gene Expression Master Mix (IDT #1055772) and a Bio-Rad CFX96 Touch Real-Time System. Data were analyzed using the 2^-ΔΔC^_T_ method with all values normalized to housekeeping genes then set relative to Day 2 gene expression levels.

### 2.7 SDS polyacrylamide gel electrophoresis and immunoblotting

All samples were run on 16% Tris-Tricine polyacrylamide gels using standard tris-tricine buffer systems as detailed in (25). Proteins were electrophoretically transferred to PVDF membranes then blocked with 5% milk in TBST for one hour. Primary antibodies were diluted at varying concentrations (1:1000 to 1:5000) in solutions of 2% to 5% fat-free milk/TBST depending on each antibody and incubated with membranes overnight at 4°C. Membranes were then washed for 15 minutes using TBST on a rotary shaker four times. Then membranes were incubated with secondary antibodies, diluted in 2% milk/TBST at varying concentrations (1:10,000 to 1:30,000) for two hours at room temperature or overnight at 4°C or two hours at room temperature or overnight at 4°C. Blots were imaged using film and ECL Prime Western Blot Detection Reagent (Amersham #RPN2322).

### 2.8 Serine palmitoyltransferase (SPT) activity in whole cerebral hemispheres

As a direct measure of SPT activity, we utilized a high performance liquid chromatography/mass spectrometry method to quantitatively measure 3-KDS (3-ketodihydrosphingosine) generated in membranes from postnatal rat brains (adapted from (26)). Briefly, 75μg of total membrane protein was incubated at 37°C for 10 minutes in 180μL of reaction mix (100mM HEPES, 5mM DTT, 5mM deuterated L-serine (3,3-D2), 50μM pyridoxyl 5’-phosphate, 100μM palmitoyl-CoA) using 13−100mm screw-cap Kimax glass tubes. Reactions were quenched with 2mL 0.5N NH_4_OH then immediately placed in −20°C freezer until lipid extraction could be completed. For extraction, 250 pmol of d17:0 dihydrosphinganine was added as an internal standard then lipids were extracted by adding 2.5mL chloroform/methanol (2:1, v/v) and vortexing vigorously. Samples were centrifuged at 4,000x*g* for 5 minutes after which the upper aqueous phase was removed. The lower organic phase was then washed twice with 5mL water. After washing, the organic phase was dried under nitrogen gas then reconstituted in 500μL mobile phase. Fifty microliters were subjected to HPLC-ESI-MS/MS analysis to measure the incorporation of deuterated L-serine (3,3-D2) into 3-KDS. Lipid analysis was performed using a Shimadzu Nexera LC-30 AD binary pump system coupled to a SIL-30AC autoinjector and DGU20A5R degasser coupled to an AB Sciex 5500 quadrupole/linear ion trap (QTrap) (SCIEX Framingham, MA) operating in a triple quadrupole mode was used. Q1 and Q3 was set to pass molecularly distinctive precursor and product ions (or a scan across multiple m/z in Q1 or Q3), using N2 to collisionally induce dissociations in Q2 (which was offset from Q1 by 30-120 eV); the ion source temperature was set to 500 °C. The lipids were separated by reverse phase LC using a Supelco 2.1 (i.d.) x 50 mm Ascentis Express C18 column (Sigma, St. Louis, MO) and a binary solvent system at a flow rate of 0.5mL/min with a column oven set to 35°C. Prior to injection of the sample, the column was equilibrated for 0.5 min with a solvent mixture of 95% mobile phase A1 (CH_3_OH/H_2_O/HCOOH, 58/41/1, v/v/v, with 5mM ammonium formate) and 5% mobile phase B1 (CH_3_OH/HCOOH, 99/1, v/v, with 5mM ammonium formate), and after sample injection (typically 40 mL), the A1/B1 ratio was maintained at 95/5 for 2.25 min, followed by a linear gradient to 100% B1 over 1.5 min, which was held at 100% B1 for 5.5 min, followed by a 0.5 min gradient return to 95/5 A1/B1. The column was re-equilibrated with 95:5 A1/B1 for 0.5 min before the next run. SPT activity is reported as pmoles of 3-KDS-D2 per mg total protein.

### 2.9 Lipidomics analysis of sphingoid bases in cerebral hemispheres Extraction of Sphingolipids

Samples were collected into 13 × 100 mm borosilicate tubes with a Teflon-lined cap (catalog #60827-453, VWR, West Chester, PA). Then 2 mL of CH_3_OH and 1 mL of CHCl_3_ were added along with the internal standard cocktail (250 pmoles of each species dissolved in a final total volume of 10 μl of ethanol:methanol:water 7:2:1). The contents were dispersed using an ultra sonicator at room temperature for 30 s. This single phase mixture was incubated at 48°C overnight. After cooling, 150 μl of 1 M KOH in CH_3_OH was added and, after brief sonication, incubated in a shaking water bath for 2 h at 37°C to cleave potentially interfering glycerolipids. The extract was brought to neutral pH with 12 μl of glacial acetic acid, then the extract was centrifuged using a table-top centrifuge, and the supernatant was removed by a Pasteur pipette and transferred to a new tube. The extract was reduced to dryness using a Speed Vac. The dried residue was reconstituted in 0.5 ml of the starting mobile phase solvent for LC-MS/MS analysis, sonicated for approximately 15 sec, and then centrifuged for 5 min in a tabletop centrifuge before transfer of the clear supernatant to the autoinjector vial for analysis.

### LC-MS/MS of sphingoid bases, sphingoid base 1-phosphates, and complex sphingolipids

These compounds were separated by reverse phase LC using a Supelco 2.1 (i.d.) x 50 mm Ascentis Express C18 column (Sigma, St. Louis, MO) and a binary solvent system at a flow rate of 0.5 mL/min with a column oven set to 35°C. Prior to injection of the sample, the column was equilibrated for 0.5 min with a solvent mixture of 95% mobile phase A1 (CH_3_OH/H_2_O/HCOOH, 58/41/1, v/v/v, with 5 mM ammonium formate) and 5% mobile phase B1 (CH_3_OH/HCOOH, 99/1, v/v, with 5 mM ammonium formate), and after sample injection (typically 40 μL), the A1/B1 ratio was maintained at 95/5 for 2.25 min, followed by a linear gradient to 100% B1 over 1.5 min, which was held at 100% B1 for 5.5 min, followed by a 0.5 min gradient return to 95/5 A1/B1. The column was re-equilibrated with 95:5 A1/B1 for 0.5 min before the next run.

### Analysis of Glucosylceramide and Galactosylceramide

Because glucosylceramide (GlcCer) and galactosylceramide (GalCer) coelute by the above method, biological samples that contain both can be analyzed by a separate method. Dried samples were re-dissolved in CH_3_CN/CH_3_OH/H_3_CCOOH (97:2:1) (v,v,v) with 5 mM ammonium acetate. The LC-Si column (Supelco 2.1 x 250 mm LC-Si) was pre-equilibrated with CH_3_CN/CH_3_OH/H_3_CCOOH (97:2:1) (v,v,v) with 5 mM ammonium acetate for 1.0 min at 1.5 mL per min, sample was injected, and the column was isocratically eluted for 8 min. GlcCer eluted at 2.56 min and GalCer at 3.12 min using this isocratic normal phase system; however, column age and previous sample load can influence the retention times of HexCer’s by this method. Periodic retention time confirmation with internal standards allowed the monitoring of column stability and subsequent effectiveness.

### Analysis of Sulfatide

In experiments where sulfatide analysis was required, 250 pmoles of the sulfatide internal standard (d18:1/C12:0 sulfatide, Avanti Polar Lipids (Alabaster, AL)) was added with the other internal standards during extraction. Sulfatides were separated by reverse phase LC using a Supelco 2.1 (i.d.) x 50 mm Ascentis Express C18 column (Sigma, St. Louis, MO) and a binary solvent system at a flow rate of 0.7 mL/min with a column oven set to 60°C. Prior to injection of the sample, the column was equilibrated for 0.5 min with a solvent mixture of 99% mobile phase A1 (CH_3_OH/H_2_O/HCOOH, 65/34/1, v/v/v,) and 1% mobile phase B1 (CH_3_OH/HCOOH, 99/1, v/v,), and after sample injection (typically 10 μL), the A1/B1 ratio was maintained at 99/1 for 3.0 min, followed by a linear gradient to 100% B1 over 2.25 min, which was held at 100% B1 for 4.5 min, followed by a 0.5 min gradient return to 99/1 A1/B1. The column was re-equilibrated with 99:1 A1/B1 for 0.5 min before the next run.

Sulfatides were analyzed in negative ion mode, using an m/z 240.9 product ion (indicative of the sulfated galactose) for quantitation. Previously sulfatide standards had been used to confirm LC retention in addition to N-acyl fatty acid product ions.

## Results

To form the basis for understanding the regulation of sphingolipid metabolism during myelination in the central nervous system we utilized the well-characterized myelination program in the neonatal rat. Rat pups were sacrificed at intervals from postnatal day 2 to postnatal day 30. To avoid litter-specific variation, pups were selected from several litters at each time-point. To assess the timing of myelination in this model we measured the protein and RNA levels of three major myelin proteins, myelin oligodendrocyte protein (MOG), myelin basic protein (MBP), and myelin proteolipid protein (PLP) (Figure 1). The mRNA expression for these three proteins in the rat brain becomes readily detectable at about 9-10 days after birth, and continues to rapidly increase until around 20 days of age. This pattern is closely followed by the rapid and sharp accumulation of the respective PLP, MOG, and major splicing isoforms of MBP protein levels. These data are consistent with previous reports that myelination in the rat brain is initiated at around postnatal day 9 and continues to increase until day 20, subsequently decreasing to reach basal levels by one month of age, when the initial round of myelination is essentially complete (27,28).

**Figure 1.**
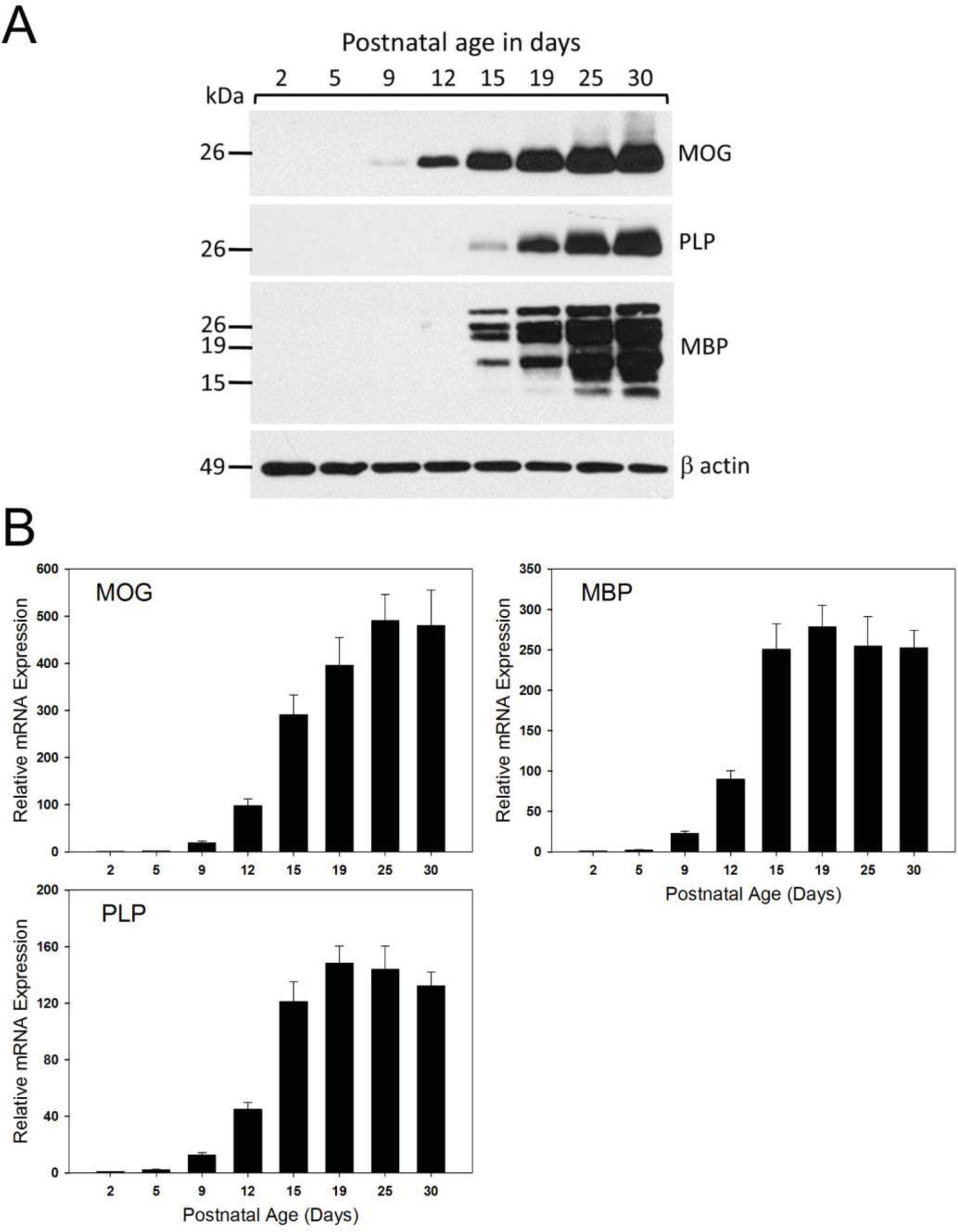
Key proteins associated with myelination increase with age in the developing rat brain. **A)** Western blot analysis of total membranes isolated from the brains of postnatal rats of various ages as indicated in each panel. Blots were probed with primary antibodies for the myelin-associated proteins MOG, PLP, MBP and Δ-actin as a loading control. **B)** Total RNA was isolated from the brains of postnatal rats of various ages as indicated. RT-qPCR analysis was performed and mRNA expression levels of key genes associated with myelination were quantitated. Data have been normalized to GAPDH expression levels then set relative to Day 2. Shown are mean plus and minus SEM, *n*=5 animals per age group. <u>MOG</u> (myelin oligodendrocyte glycoprotein); <u>PLP</u> (proteolipid protein); <u>MBP</u> (myelin basic protein); <u>HPRT</u> (hypoxanthine phosphoribosyltransferase).

We first performed a profiling of sphingolipids in total brain during this period of myelination. In interpreting this data it is important to recognize that in addition to the initiation of myelination, the first two postnatal weeks also include synaptogenesis and increases in neuronal, astroglial, and microglial cells and these will contribute to the overall sphingolipid profile. Nevertheless, galactocerebrosides and their sulfated derivative, sulfatide, are considered to be specific oligodendroglial and myelin components. Our analysis included only sphingolipids containing non-hydroxylated fatty acids. However it should be emphasized that the contribution of hydroxyl sphingolipids, particularly galactosylceramide, to myelin production and function is important and has been the attracted considerable attention (29,30).

### Levels of the sulfatide, monohexosylceramides, galactosylceramide sulfotransferase, and ceramide galactosyltransferase in the developing brain begin to elevate at postnatal day 5

The sulfated sphingolipid, 3-O-sulfogalactosylceramide, referred to as sulfatide, is a distinct component of myelin. Using mass spectrometry we measured levels of sulfatide in these samples and found that they follow the levels of expression of the myelin-specific proteins (Figure 2, Panel A). Levels of sulfatide start rising above baseline at day 5, then rise steadily until day 25, where they plateau. This elevation is mirrored by increased expression of Gal3st, the enzyme that produces sulfatide by sulfation of galactosylceramide (Figure 2, Panel B). We also measured the combined levels of the mono-glucosylated ceramide species (collective known as monohexosylceramides or cerebrosides) glucosylceramide and galactosylceramide. Monohexosylceramides rise above background at day 9 and then steadily increase with a plateau at day 25. This is similar, but slightly slower than the timing of sulfatide production. Expression of the ceramide galactosyltransferase (Ugt8) rises precipitously from day 9 to day 15, where it plateaus. This mirrors early measurement of activity and expression levels of this enzyme (31,32). This is consistent with an increased production of galactosylceramide as a component of myelin as well as to fuel the production of sulfatide.

**Figure 2.**
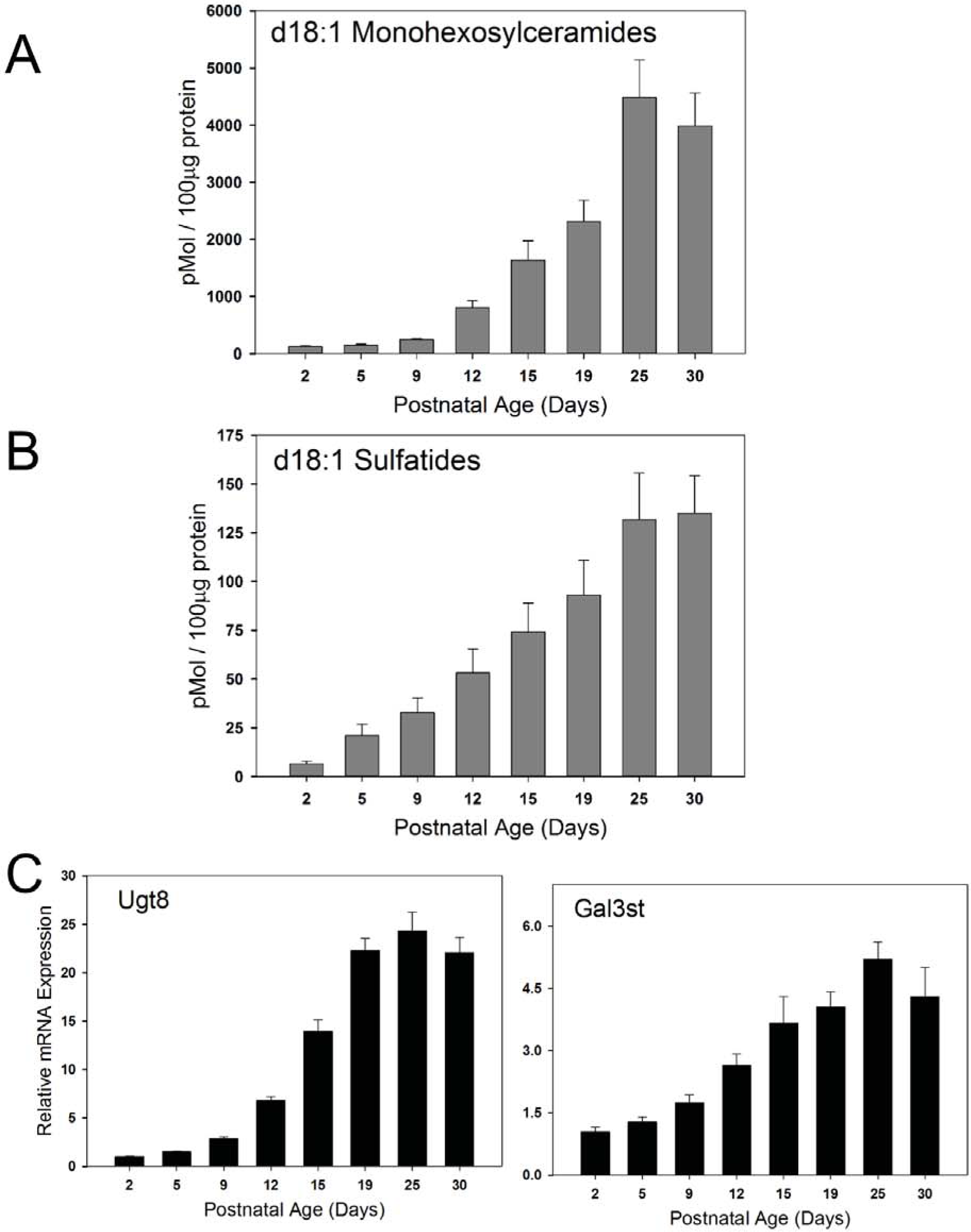
Mass levels of key sphingolipids and corresponding sphingolipid metabolic enzymes associated with myelination increase in the postnatal rat brain. Total membranes were isolated from the brains of postnatal rats of various ages as indicated. Steady state levels of d18:1:Monohexosylceramides **(A)** and d18:1:Sulfatides **(B)** were quantitated using LC-MS/MS analysis. Data are graphed as pMol lipid per 100μg total protein. **C)** Total RNA was isolated from the brains of postnatal rats of various ages as indicated. RT-qPCR analysis was performed and mRNA expression levels of key enzymes responsible for the biosynthesis of monohexosylceramides and sulfatides were quantitated. Data have been normalized to GAPDH expression levels then set relative to Day 2. Presented as mean (+/-) SEM, *n*=5 animals per age group. <u>Ugt8</u> (UDP-Galactose-Ceramide Galactosyltransferase 8); <u>Gal3st</u> (Galactose-3-O-sulfotransferase).

### Levels of ceramides and sphingomyelins rise moderately in the myelinating brain. Levels of S1P and dihydroS1P rise late in the period of myelination

Ceramide is an obligate intermediate in the production of the sphingolipid components of myelin, but at elevated levels is pro-apoptotic (33). It is therefore not surprising that the levels of ceramide stay relatively constant throughout the course of myelination (Figure 3, Panel A) rising gradually to a plateau level at postnatal day 19 that is approximately 2 fold over that at postnatal day 2. This is somewhat similar to other recent observations by Dasgupta and Ray using slightly different techniques (34). Levels of ceramide are approximately an order of magnitude lower than that of the monohexosylceramides. Levels of sphingomyelin, a component of the myelin membrane, are also relatively constant, rising less than 2 fold in total brain lipids during the course of myelination (Figure 3, Panel B). This is somewhat surprising considering that sphingomyelin is a significant component of myelin. However the mass levels of sphingomyelin are high, so even moderate increases represent a significant contribution.

**Figure 3.**
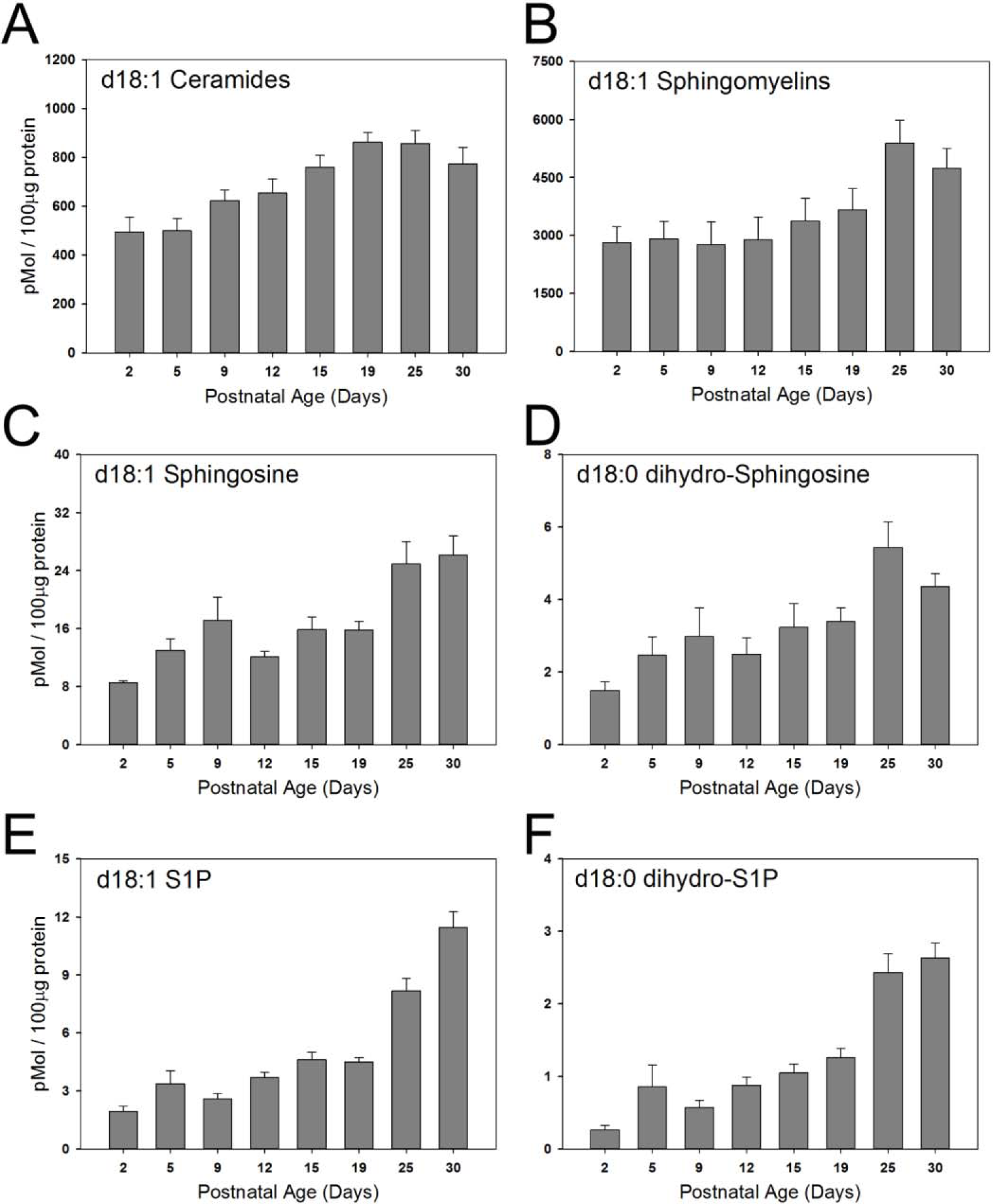
Steady state levels of d18:1 sphingolipids in the membranes of the post-natal rat brain. Total membranes were isolated from the brains of postnatal rats of various ages as indicated in each panel. Steady state levels of total d18:1:Ceramides **(A)**, d18:1:Sphingomyelins **(B)**, d18:1 Sphingosine **(C)**, d18:1 dihydro-Sphingosine **(D)**, d18:1 S1P **(E)**, and d18:1 dihydro-S1P **(F)** were quantitated using LC-MS/MS analysis. Data are graphed as pMol lipid per 100μg total protein, Mean (+/-) SEM, *n*=5 animals per age group.

Sphingosine and dihydrosphingosine are upstream metabolites in the sphingolipid pathway. Sphingosine is a product of ceramide hydrolysis by various ceramidases, whereas dihydrosphingosine is produced early in the biosynthetic pathway prior to ceramide synthesis. Levels of sphingosine rise gradually during postnatal development (Figure 3, Panel C) to levels at postnatal day 25 that are approximately 3-fold over that at postnatal day 2. This may reflect both the elevation of ceramides as well as the activity of ceramidases. Levels of dihydrosphingosine also rise during the period of myelination, but are elevated more dramatically late in the process (Figure 3, Panel D). Although this may be a reflection of elevated *de novo* sphingolipid biosynthesis, it is important to recognize that these are steady-state measurements, so levels are the result of both production and consumption of metabolic intermediates. This flux is consistent with the observation that the levels of the sphingosines are hundreds of fold lower than that of the downstream sphingolipid metabolites. The phosphorylated derivatives of the sphingosines, dihydrosphingosine-1-phosphate and sphingosine-1-phosphate are well-studied extra- and intracellular signaling molecules and metabolic intermediates (reviewed in (21)). Interestingly levels of both of these molecules rise steadily after birth, but, in contrast to the specific myelin lipids, do not plateau at postnatal day 25, but continue to rise (Figure 3, Panels E and F). The time-course of this elevation, which continues past the peak time of myelination, may indicate that these lipids are important players in neurodevelopmental processes and in the mature rat brain (reviewed in (35)). In support of the notion that the phosphorylated sphingosines may not have a crucial role in the process of myelination, we do not observe an increase in the phosphorylated sphingosines in isolated oligodendrocytes during this period (Supplementary Figure 1).

### Levels of the major sphingolipids in oligodendrocytes during myelination

The measurements presented above are derived from analysis of cerebral hemispheres during the postnatal time window that includes the active period of myelination. While some of the lipids and enzymes measured, such as sulfatides and Gal3T clearly reflect processes occurring in the myelin-producing oligodendrocytes, other measurements may include contributions from neurons, astrocytes, and microglia, which are all proliferating and differentiating during this period (reviewed in (36)). To address this issue we isolated oligodendrocytes at days 2, 8, and 16 after birth, to capture the period of peak myelination. To ensure sufficient material for valid measurements these data are derived from pooled oligodendrocytes isolated from the brains of 20 rat pups. In oligodendrocytes levels of monohexosylceramides are relatively low in cells isolated from 2 and 8-day old pups, but exhibit a dramatic elevation in cells isolated 16 days after birth (Figure 4). Here we fractionated the monohexosylceramides into the two major components, glucosylceramide and galactosylceramide. These show similar temporal patterns, with glucosylceramide composing the major species. This is interesting and surprising since glucosylceramide is essentially absent from myelin (37). Sulfatides rise in a similar pattern as the cerebrosides. In isolated oligodendrocytes, as in total brain, the levels of ceramides rise slowly during this time period. However in contrast to total brain, the levels of sphingomyelin undergo a significant elevation in oligodendrocytes as myelination proceeds. It is important to note that during the isolation of the oligodendrocytes it is likely that fragile processes are sheared and lost. Consequently, it is likely that these measurements reflect levels of lipids predominantly in the cell body.

**Figure 4.**
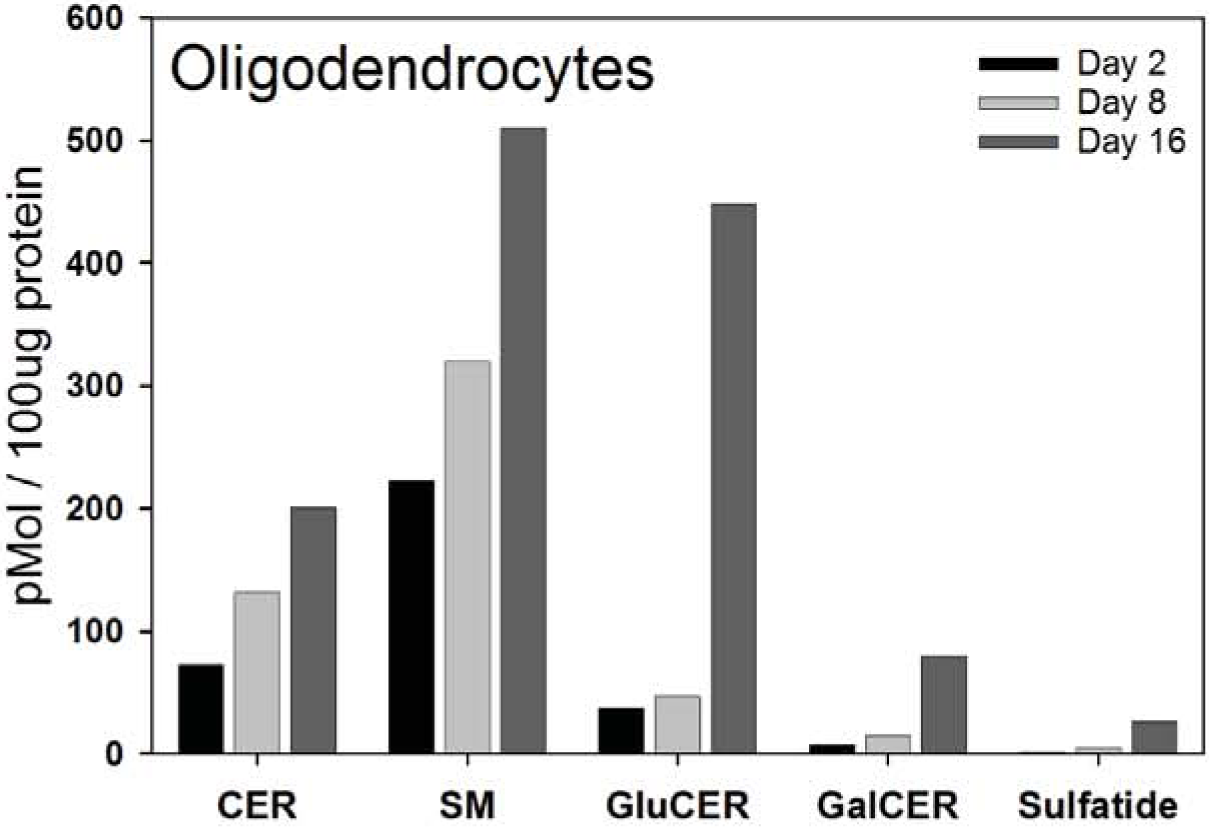
Steady state levels of total d18:1 sphingoid base lipids were measured in both the oligodendrocytes isolated from the brains of postnatal rats of various ages. LC-MS/MS was used to quantitate the mass levels of total d18:1 ceramides (CER), d18:1 sphingomyelins (SM), d18:1 glucosylceramides (GluCER) d18:1 galactosylceramides (GalCER), and d18:1 sulfatides (Sulfatide) in both the primary oligodendrocytes isolated from the brain tissue of postnatal rats using the Percoll gradient method as described in *Materials and Methods Section 2.3*. Data are graphed as pMol lipid per 100μg total protein. Samples are from pooled oligodendrocytes derived from 20 rat pups.

### Activity and protein levels of serine palmitoyltransferase and associated regulators in total cerebral hemispheres

Serine palmitoyltransferase (SPT) is the initiating and rate-limiting step in sphingolipid biosynthesis (reviewed in (13)). The core enzymatic unit of SPT is a heterodimer consisting most often of SPTLC1and SPTLC2. Both subunits are required for activity although the active site is contained within SPTLC2. Less commonly SPTLC3 (also referred to as SPTLC2b) replaces SPTLC2 in the complex. In addition, there are two smaller subunits, ssSPTa and –b which can profoundly affect SPT activity and substrate specificity (14). While palmitoyl (16:0)-CoA is the preferred substrate for SPTLC1/2/ssSPTa, other combinations with SPTLC3 and/or ssSPTa or –b result in a complex that can accommodate myristoyl (14:0)-CoA and/or stearoyl (18:0)-CoA (14). SPT activity is homeostatically regulated in response to changes in cellular sphingolipid levels. This is mediated by small membrane proteins, the ORMDLs, of which there are three isoforms in mammalian cells (Reviewed in (17)). Given the extent of sphingolipid biosynthesis required to produce myelin it is expected that the SPT complex is tightly regulated during the early period of myelination. The importance of that regulation is supported by studies in mice conditionally overexpressing the core SPT complex in myelin-producing cells, which demonstrate severe myelination defects (38).

SPT activity was measured in total brain membranes using a sensitive and specific mass spectrometry-based assay (26). Interestingly, activity increased from day 2 to day 5 after birth and declined thereafter (Figure 5, Panel A). This decline is surprising considering the increase in sphingolipid synthesis that is occurring during this time-course. Measurement of the protein levels of the major SPT subunits revealed that levels of SPTLC2 roughly mirror the levels of SPT activity, while levels of SPTLC1 were relatively constant (Figure 5, Panel B). This is consistent with the mRNA levels of these proteins as measured by real-time PCR (Figure 6, Panels A and B). SPTLC3 can replace SPTLC2 in the core SPT complex. Intriguingly, both protein (Figure 5, Panel B) and mRNA (Figure 6, Panel C) levels of SLPTLC3 increase to a peak at days 12 and 15, suggesting a change in the composition of the SPT complex during this critical phase of myelination. Also interesting is the complementary change in levels of the small SPT subunits ssSPTa and –b. ssSPTa mRNA levels decrease dramatically during the phase of myelination (Figure 6, Panels D and E), while levels of ssSPTb increase. These subunits affect both the activity and substrate specificity of SPT, and how this might impact SPT activity during myelination is outlined in the Discussion. The three ORMDL isoforms cannot be distinguished by any available antibodies. The total level of ORMDL protein, which appears relatively constant over the time-course measured here (Figure 5, Panel B), therefore reflects the combined levels of these isoforms. However, probing isoform-specific expression at the RNA level (Figure 7) reveals that there is a dramatic decrease in the level of ORMDL1 as myelination progresses. ORMDL3 expression declines in a somewhat more modest way, while ORMDL2 levels remain constant. It should be kept in mind that activity, protein, and mRNA measurements reflect changes in all cell types in the brain. As described below a significantly different picture emerges from measurements made in isolated oligodendrocytes.

**Figure 5.**
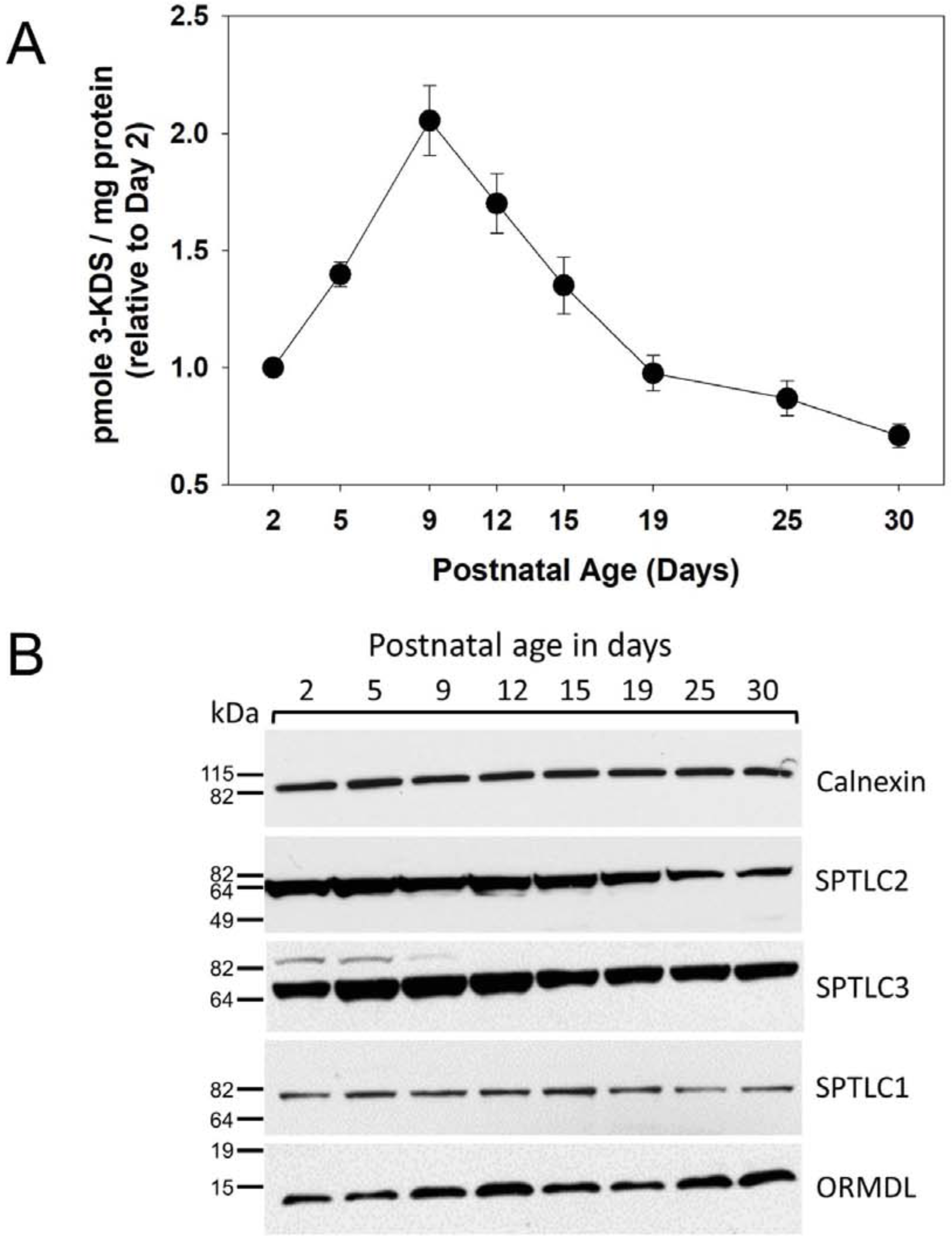
Dynamic changes in the activity and subunit composition of serine palmitoyltransferase in the early post-natal rat brain. **A)** Serine palmitoyltransferase activity was measured using an HPLC-ESI-MS/MS method combined with deuterated serine to quantitatively measure 3-KDS (3-Ketodihydrosphingosine) generated in membranes from postnatal rat brains. Data are graphed as pMol 3-KDS lipid per mg total protein, Mean (+/-) SEM, *n*=5 animals per age group. **B)** Western blot analysis of total membranes isolated from the brains of postnatal rats of various ages as indicated in each panel. Blots were probed with primary antibodies for the large subunits of serine palmitoyltransferase (SPTLC1-3), total ORMDL protein and calnexin.

**Figure 6.**
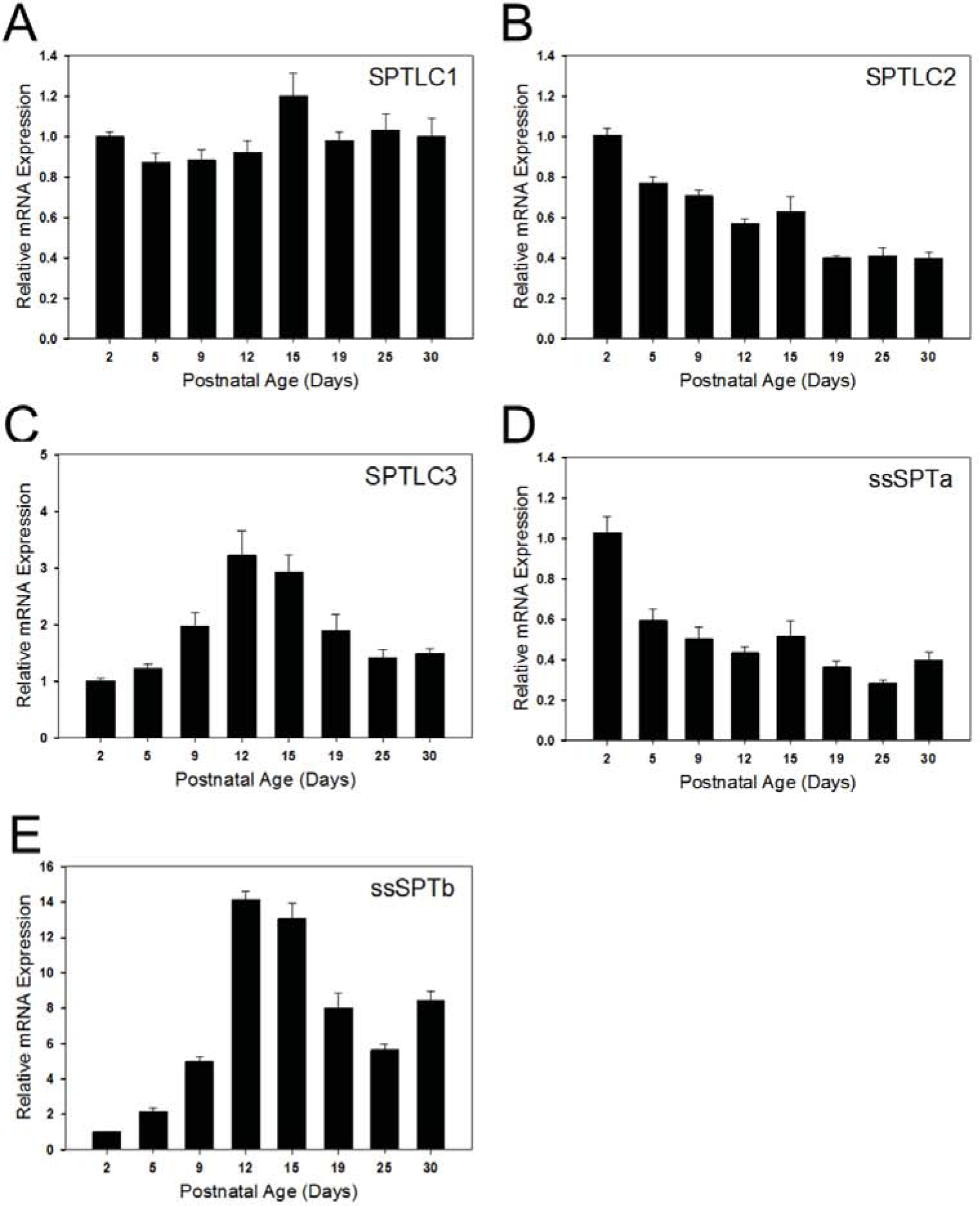
mRNA expression levels of the serine palmitoyltransferase large and small subunits are dynamically regulated in the developing rat brain. Total RNA was isolated from the brains of postnatal rats of various ages as indicated in each panel. RT-qPCR analysis was performed and mRNA expression levels of the large subunits, SPTLC1 **(A)**, SPTLC2 **(B)**, and SPTLC3 **(C)**, as well as the small subunits, ssSPTa **(D)** and ssSPTb **(E)** were quantitated. Data have been normalized to GAPDH expression levels then set relative to Day 2. Presented as mean (+/-) SEM, *n*=5 animals per age group.

**Figure 7.**
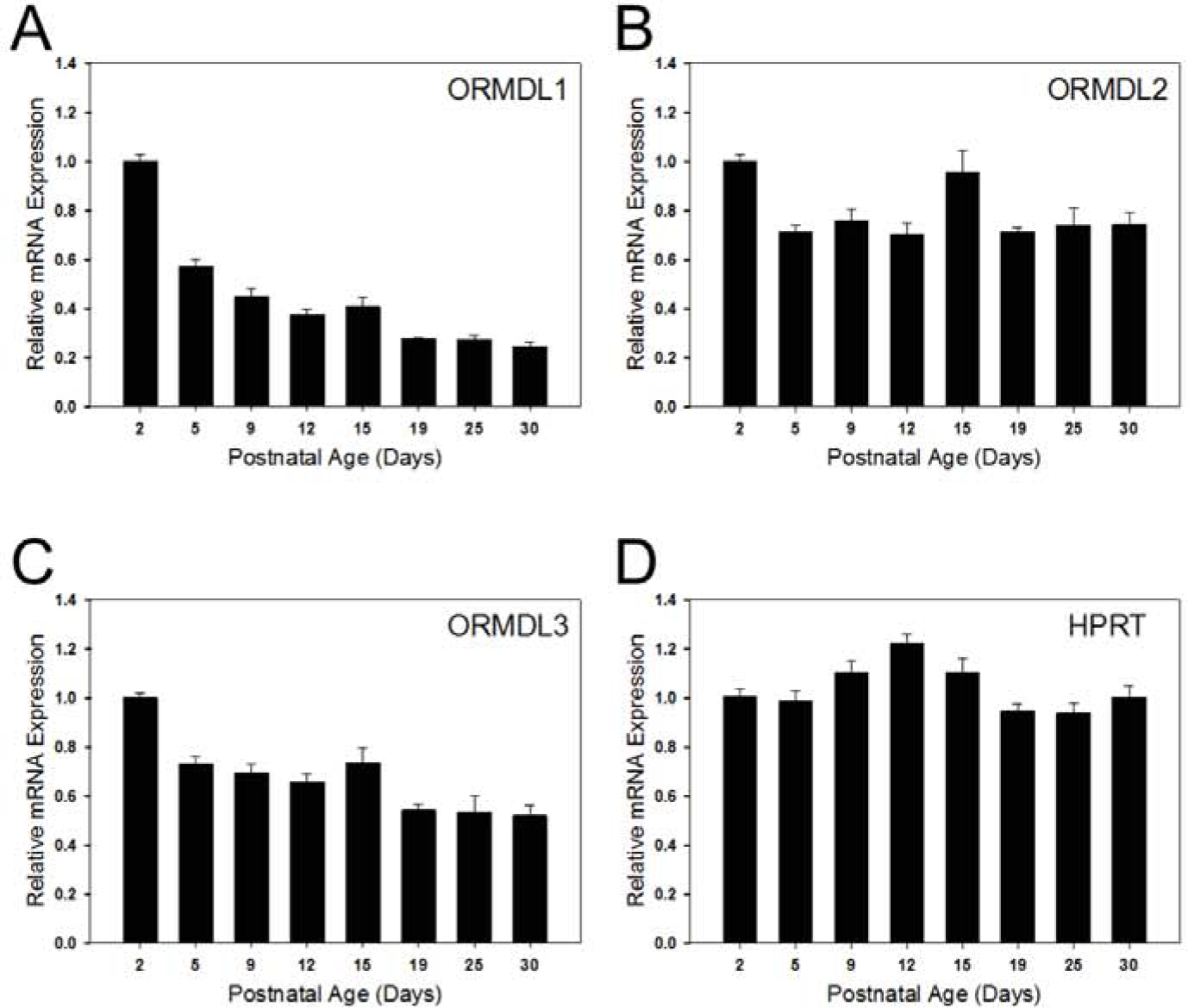
mRNA expression levels of the three ORMDL isoforms are dynamically regulated in the developing rat brain. Total RNA was isolated from the brains of postnatal rats of various ages as indicated in each panel. RT-qPCR analysis was performed and mRNA expression levels of ORMDL1 **(A)**, ORMDL2 **(B)**, and ORMDL3 **(C)**, as well as the housekeeping gene, HPRT **(D)** were quantitated. Data have been normalized to GAPDH expression levels then set relative to Day 2. Shown are mean (+/-) SEM, *n*=5 animals per age group.

### Levels of sphingolipid metabolic enzymes, SPT subunits, and ORMDL isoforms in isolated oligodendrocytes

As expected, in isolated oligodendrocytes expression of the genes encoding the myelination marker proteins MBP and MOG, as measured at the protein and mRNA levels, are virtually absent in cells isolated from 2 day-old pups, is greatly elevated as myelination progresses from 8 to 16 days of age (Figure 8, Panels A and D.) Expression of Ugt8, the gene encoding the enzyme which produces galactosylceramide, is virtually undetectable in oligodendrocyte progenitors from 2 day-old pups but dramatically increases as more cells transition from pre-oligodendrocytes into differentiated cells at the beginning of the rapid period of myelination at around postnatal day 8 (Figure 8, Panel D.) Ugt8 mRNA remains at similarly elevated levels in cells obtained at postnatal day 16, when the majority of isolated oligodendrocytes are almost exclusively mature oligodendrocytes capable of myelination. In contrast, mRNA levels of Gal3T, which produces sulfatide from galactosylceramide, are still low in cells from 8-day-old pups; but increase by over 500-fold by the time cells are already fully differentiated. These data confirm the expectation that the majority of the changes in galactosylceramide and sulfatide content in the whole brain during myelination is driven by oligodendrocyte activity (39).

**Figure 8.**
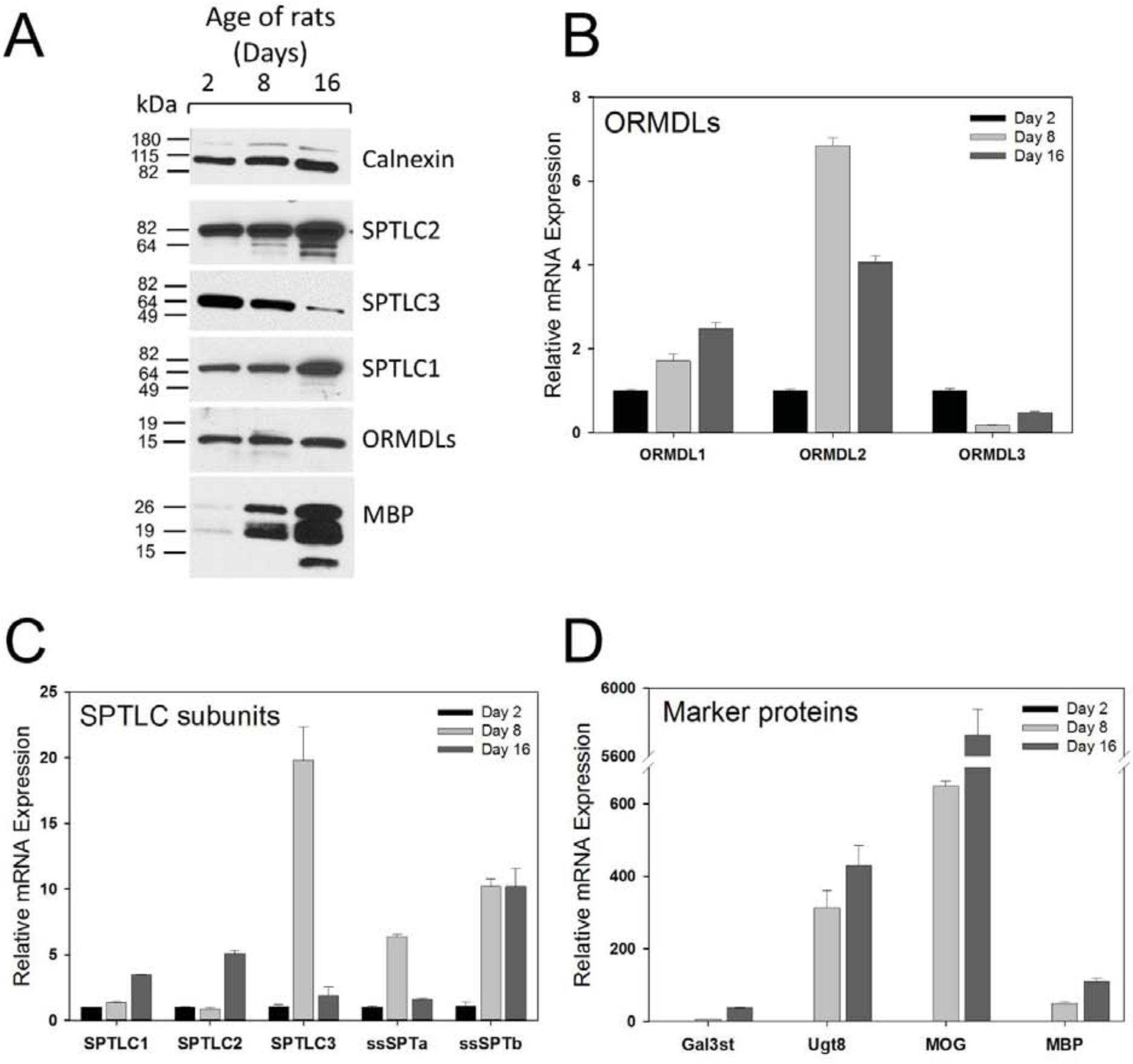
Serine palmitoyltransferase and ORMDL proteins are temporally regulated in oligodendrocytes isolated from the brains of postnatal rats of various ages. Primary oligodendrocytes were isolated by Percoll gradient purification as detailed in *Materials and Methods*. After isolation, half of the cells were homogenized and used for Western blot analysis of SPT subunits and MBP as depicted in **(A)** Calnexin is included as a loading control. The remaining cells were used for total RNA isolation and real-time qPCR analysis to quantitate the mRNA expression levels of the ORMDL isoforms **(B)**, the serine palmitoyltransferase subunits **(C)**, as well as oligodendrocyte-specific genes **(D)**. mRNA expression data were normalized to GAPDH expression levels then set relative to Day 2. Shown is mean (+/-) SEM of triplicate technical replicates. Shown is one of two determinations which yielded similar results.

To determine whether changes in the SPT complex in the intact brain reflect progression of the oligodendrocyte development and myelination program we measured the expression levels of SPT subunits in isolated oligodendrocytes (Figure 8). Levels of SPTLC1 and 2 progressively increased at both the protein and mRNA levels in oligodendrocytes as the cells differentiate from immature oligodendrocyte progenitors (day 2) into mature myelinating oligodendrocytes (day 16) (Figure 8, Panels A and B). This contrasts to a degree to the measurements in whole brain, where levels of SPTLC1 were constant and SPTLC2 declined over the same period (Figures 5 and 6). These data indicate that cell types other than oligodendrocytes influence the measurement of these genes in the intact brain. The increase in SPT in oligodendrocytes is consistent with the increased need to supply the sphingolipids required for myelination. Interestingly, protein levels of SPTLC3 decline during this period (Figure 8, Panel A) although there is a dramatic increase in SPTLC3 message levels at day 8, (Figure 8, Panel C). The discrepancy between these two measurements has yet to be resolved. The day 8 elevation of SPTLC3 somewhat parallels the measurements in whole brain, although in the brain the levels of SPTLC3 continue to elevate through day 15 and beyond. The expression of the small subunits also reveal oligodendrocyte-specific changes in the composition of the SPT complex (Figure 8, Panel C). In oligodendrocytes the levels of ssSPTa are near baseline at day 16. This contrasts to the decline of ssSPTa over the entire time course measured in whole brain (Figure 6). ssSPTb message is markedly elevated at day 8 and is maintained at this level at day 16. This is qualitatively similar to what is observed in the whole brain samples (Figure 5). The overall protein levels of the ORMDLs do not dramatically change during this period (Figure 8, Panel A), however measurement of mRNA of the individual isoforms indicate that the relative expression of the three isoforms changes significantly (Figure 8, Panel B). Expression of ORMDL1 and −2 increase during myelination while levels of ORMDL3 decline. These data indicate that the overall levels of the SPT complex increase during the peak of myelination, but that the composition of the complex changes both with respect to the major and minor subunits, driving changes both in the levels and molecular species of sphingolipids produced.

### Levels of d16:1 and d20:1 sphingoid backbones in oligodendrocytes during myelination

The SPTLC1/2 dimer in combination with ssSPTa strongly prefers palmitoyl(16:0)-CoA as a substrate, and therefore generates sphingolipids containing the d18 sphingoid base. However other trimer combinations exhibit a variety of acyl-CoA substrate preferences (14). For example SPTLC1/3/ssSPTa can use myristic (14:0)-CoA equivalently to palmitoyl-CoA, yielding d16 bases and SPTLc1/3/ssSPTb can utilize either myristoyl- or stearoyl (18:0) CoA’s. Therefore the composition of the SPT complex would be expected to determine the exact molecular species of sphingoid bases produced. To examine the impact of the changes in SPT complex composition on the utilization of either myristoyl- and/or stearoly CoA we examined the levels of sphingolipids incorporating d16:1 and d20:1 sphingoid bases in isolated oligodendrocytes during the period of peak myelination (Figure 9). Because overall levels of sphingolipids change during the time-course examined (Figure 4) we have depicted the levels of these lipids as a percentage of the major d18:1 containing sphingolipids. d16:1 sphingolipids were barely detectable in ceramides and monohexosylceramides, but constitute a significant fraction of sphingomyelins, although there is only a moderate change in these levels over time (Figure 9, Panel A). In contrast, d20:1 sphingolipids constitute a significant fraction of all sphingolipids measured (Figure 9, Panel B). Levels of d20:1 ceramide and sphingomyelin rise considerably during this period, eventually constituting almost 8% of the totals of these lipids. The levels of d20:1 monohexosylceramides rise to the same level by day 8, but then decline to approximately 3% of the total.

**Figure 9.**
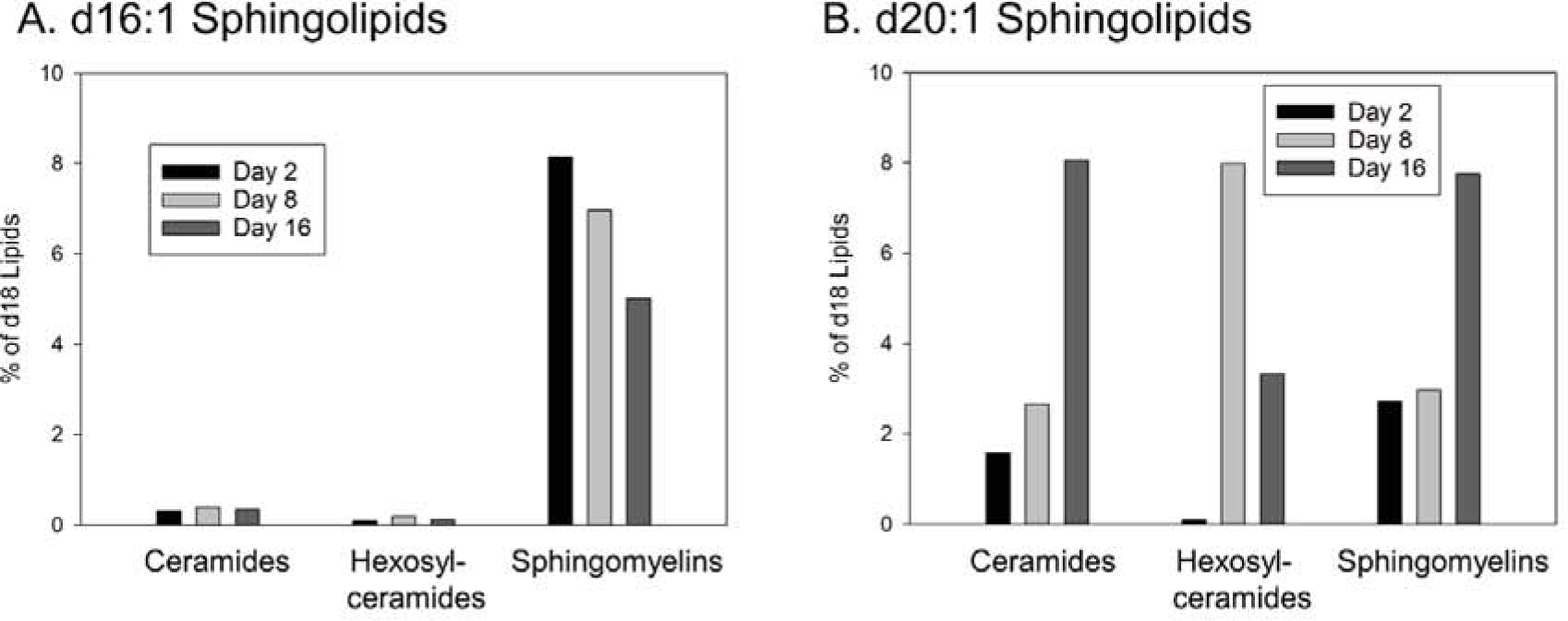
The incorporation of d16:1 and d20:1 sphingoid bases into sphingolipids in oligodendrocytes during myelination. Oligodendrocytes were isolated from the brains of rat pups at days 2, 8, and 16 after birth, lipids were extracted and subjected to analysis by mass spectrometry. **(A)** Levels of d16:1 sphingoid base incorporated into ceramides, monohexosylceramides, and sphingomyelin. **(B)** Levels of d20:1 sphingoid base incorporated into ceramides, monohexosylceramides, and sphingomyelin. Data are graphed as % of each species as a total of d18:1 sphingolipid. Mass levels of these samples are presented in *Supplementary Materials.* Shown are single samples from oligodendrocytes isolated from 20 pups.

### Analysis of the molecular species of sphingolipids in the brain and oligodendrocytes during the onset of myelination demonstrate selectivity in the downstream metabolism of ceramide

The fatty acid N-acylated to ceramide, and therefore incorporated into more complex sphingolipids, can vary from 14-26 carbons and may be either saturated on monounsaturated. The biophysical and biological properties of sphingolipids is strongly affected by the N-acyl chain lengths of those lipids (40) (reviewed in (41)) and is known to affect myelination (42). We found that the fatty-acyl content of sphingolipids in total brain and isolated oligodendrocytes changes dramatically during the period of myelination (Figure 10). For clarity we show only the 16:0, 18:0, 24:0, and 24:1 molecular species. These comprise almost 80% or more of the total species under almost all circumstances. The most notable exception is sphingomyelin in oligodendrocytes, in which 18:1 comprises 30% of the total at day 16. Overall there is a decrease in the incorporation of the shorter chain species and an elevation of the longer chain species as myelination progresses. However the individual types of sphingolipids demonstrate dramatic differences in the dynamics of change and exhibit major differences in which species predominate.

**Figure 10.**
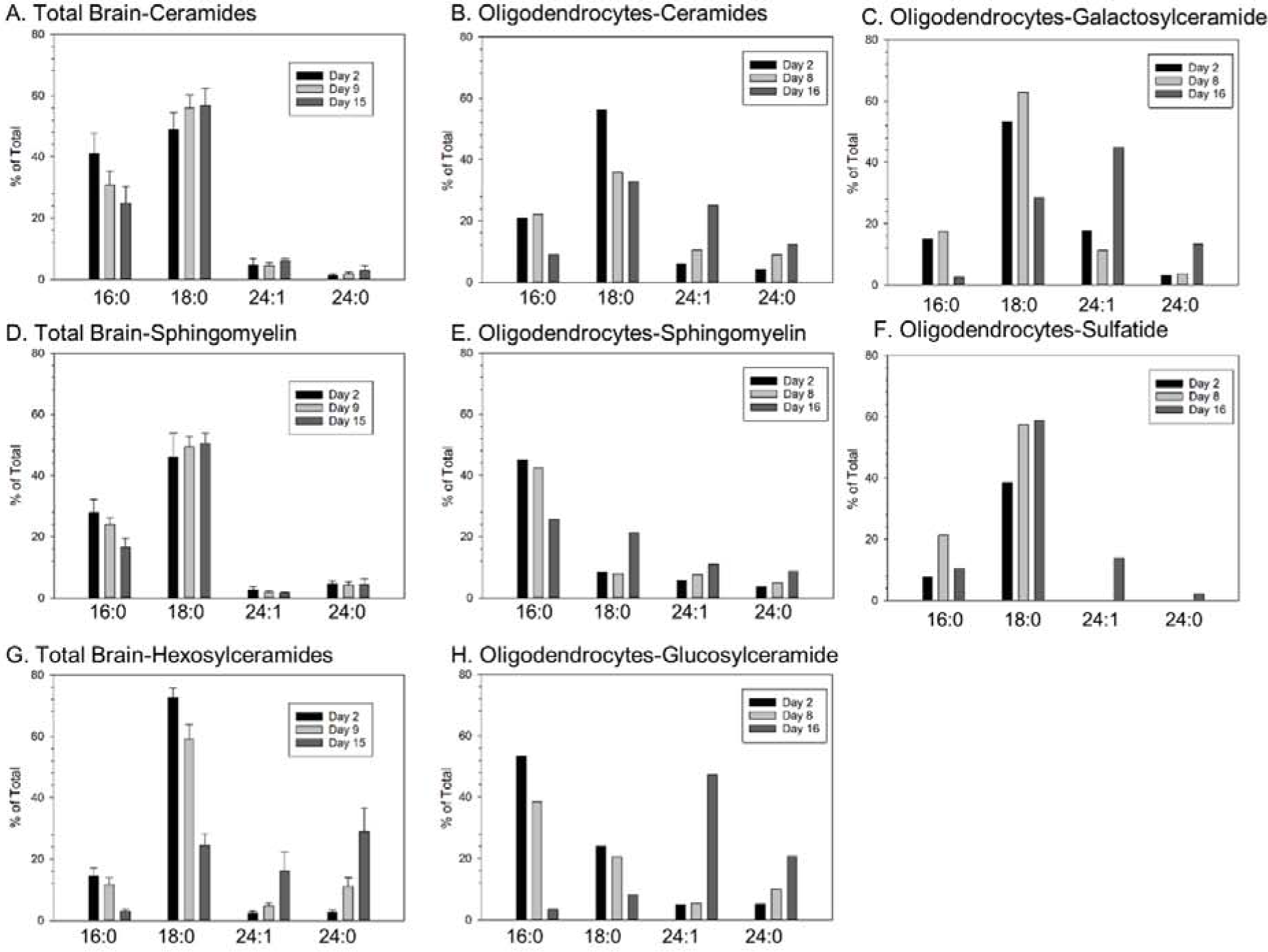
The molecular species of sphingolipids change dramatically over time and differ between total brain and isolated oligodendrocytes in the post-natal rat brain. Membranes prepared from total brain **(A, D, G)** or isolated oligodendrocytes **(B, C, E, F, H)** just prior to myelination (post-natal day 2), early in myelination (day 9 for total brain, day 8 for oligodendrocytes), and during peak myelination (day 15 in brain, day 16 in oligodendrocytes) were analyzed by mass spectrometry. For clarity shown are only the major molecular species (with some exceptions noted in Results). **(A-B)** ceramides, **(D-E)** sphingomyelins, **(G)** total hexosylceramides, **(H)** glucosylceramides, **(C)** galactosylceramides, **(F)** sulfatides. For total brain n=5, for oligodendrocytes shown are single samples from oligodendrocytes isolated from 20 pups. For brain fractions shown is mean plus and minus standard deviation.

### Ceramides in total brain and oligodendrocytes

In total brain samples initially (2 days after birth), the major molecular species in ceramide are relatively short; the combined content of 16:0 and 18:0 species constitute approximately 90% of the total (Figure 10, Panel A) and do not markedly change over time. Oligodendrocytes at day 2 have a similar distribution to that of total brain with a total of almost 80% of the ceramide species consisting of 16:0 and 18:0 (Figure 10, Panel B). As myelination progresses there is a marked shift increase towards longer ceramide species in oligodendrocytes, but not in total brain. At day 8 in oligodendrocytes the combined 16:0/18:0 content has decreased to 58% and by day 16 to 42%. In oligodendrocytes, but not total brain, the levels of the long chain ceramides, particularly 24:1 are elevated. Levels of 24:1 are approximately 6% of the total at day 2 and rise to 25% by day 16.

### Monohexosylceramides in total cerebral hemispheres and monohexosylceramides and sulfatides in oligodendrocytes

The molecular species of the monohexosylceramides are distinctly more dynamic than those of ceramide, and although the trends are similar to those in ceramide there are some distinct differences. The combined 16:0/18:0 content of monohexosylceramides in total brain declines from 87% at day 2 to 29% at day 15 (Figure 10, Panel G). In oligodendrocytes the monohexosylceramides were fractionated into glucosyl- and galactosylceramide. Glucosylceramide comprises approximately 80% of total monohexosylceramides (Figure 4). The decline of 16:0/18:0 content of oligodendrocyte glucosylceramide follows the trend of total brain monohexosylceramide but is even more severe, declining from 77% at day 2 to just 11% at day 16 (Figure 10, Panel H). Oligodendrocyte galactosylceramide differs somewhat from glucosylceramide in having lower initial levels of 16:0 and higher levels of 18:0, with a more moderate decline in the 16:0/18:0 content as myelination progresses (Figure 10, Panel C). The 24:1 species of both glucosylceramide and galactosylceramide becomes the dominant molecular species as myelination progresses. The molecular species incorporated into sulfatides are generally similar to those in the galactosylceramides (Figure 10, Panel F).

### Sphingomyelin in total cerebral hemispheres and oligodendrocytes

The molecular species in sphingomyelin are remarkably constant over time in total brain (Figure 2, Panel D) and isolated oligodendrocytes (Figure 2, Panel E). In both sources shorter chain species predominate the composition of sphingomyelin. Interestingly, in oligodendrocytes the 18:1 molecular species, otherwise a minor species, comprises close to 30% of the total sphingomyelin content at day 16. The difference in ceramide utilization between sphingomyelin and the monohexosylceramides is an indication of selective downstream ceramide metabolism (addressed in more detail in Discussion).

## Discussion

The production of myelin by oligodendrocytes in the central nervous system requires the coordinated synthesis and assembly of a specialized set of proteins and lipids (1). Myelin is highly enriched with a distinct set of lipids compared to other biological membranes and contains sphingolipids that are not found in abundance elsewhere. The molecular mechanisms that coordinate and regulate the production of individual lipid and protein components to produce the optimal composition of this critical membrane is poorly understood. This knowledge is essential for a basic understanding of the production of this highly specialized membrane and for how dysregulation could contribute to devastating demyelinating diseases.

### Levels of sphingolipids rise in total brain and isolated oligodendrocytes

Here we have focused on the sphingolipid components of myelin: the sulfatides, cerebrosides, and sphingomyelin (Reviewed in (43)). We produced a detailed time course of the levels of these lipids in total brain. The analysis of lipids in total brain during the early post-natal peak of myelination reflects both myelin production as well as the dramatic expansion of neuronal and glial cells. To focus on myelin production we also report, for the first time, analysis of sphingolipids in oligodendrocytes during this period. In parallel we measured the expression of key metabolic enzymes in this pathway, with a focus on the initiating and rate limiting step in sphingolipid biosynthesis, serine palmitoyltransferase.

As expected, the levels of sulfatides and cerebrosides, as well as the proximal enzymes that produce the sulfatides, Gal3T and UGT8, rise in both the total brain and oligodendrocyte samples as the period of myelination progresses. Similarly, the elevation of sphingomyelin is consistent with its role as a component of myelin. More interesting is the elevation of ceramide levels as myelination progresses as we find here and as noted previously (44). Ceramide is required as a precursor for the cerebrosides, sulfatides, and sphingomyelin, but elevated ceramide levels can also trigger apoptosis (45). As discussed below, there are dynamic changes in the molecular species composition of the ceramides which may moderate the pro-apoptotic effects of this sphingolipid.

### Time and sphingolipid-dependent changes in molecular species during myelination

We and others observe that as myelination progresses there is a decline in the levels of long chain (16:0 and 18:0)-containing sphingolipids (44,46) and a corresponding increase in the very long chain sphingolipids (principally 24:0 and 24:1). In our studies this is less apparent in the total brain lipids such as ceramides and sphingomyelin, but readily seen in the myelin-enriched monohexosylceramides in total brain samples as well in all sphingolipids measured in isolated oligodendrocytes. This shift towards longer molecular species parallels a decline in the expression of Cers6, which produces long-chain ceramides and an increased expression of Cers2, which produces very long chain ceramides (44,47). The functional importance of the elevated levels of very long chain sphingolipids is underscored by the severe myelination defects that result from deletion of Cers2 (42,48). Although ceramide is generally considered to be pro-apoptotic, there is emerging evidence that this is not universal for all ceramide molecular species (49-51) (reviewed in (52)). A shift of ceramide molecular species to very long chain species may explain how an increased ceramide pool can be tolerated by myelinating oligodendrocytes.

Examination of the distribution of molecular species in oligodendrocytes reveals that there is selective incorporation of ceramide species in different classes of sphingolipids. For example sphingomyelin remains relatively rich in 16:0 through the peak period of myelination (Figure 10, Panel E) while levels of this molecular species are barely detectable in the cerebrosides (Figure 10, Panels C and H). Conversely levels of 24:1 are strongly elevated in the cerebrosides as myelination proceeds but are a minor species in sphingomyelin. In galactosylceramide and sulfatide 18:0 is the major molecular species, but this is a relatively minor species in glucosylceramide. Selectivity may be introduced at the level of intercompartmental transport. Ceramide must be transported from its site of synthesis in the endoplasmic reticulum to the trans Golgi for sphingomyelin synthesis and to the medial Golgi for glucosylceramide synthesis (Reviewed in (53)). In contrast galactosylceramide synthesis occurs in the endoplasmic reticulum and so presumably does not require a transport mechanism. The ceramide transporter, CERT, required for ceramide transport to the trans Golgi for sphingomyelin synthesis, is selective for shorter chain ceramides (54,55). This may account, in part, for the enrichment of these species in sphingomyelin. The mechanism mediating transport of ceramide to the cytosolic face of the medial Golgi for glucosylceramide synthesis is still unresolved (53,56). Whether there is selectivity in this transport process remains to be determined. The close correlation between ceramide and galactosylceramide molecular species may reflect that ceramide and galactosylceramide production are both in the endoplasmic reticulum.

A particularly striking finding of our study is that glucosylceramide is a substantial component of the sphingolipid pool in myelinating oligodendrocytes. Yet myelin contains negligible levels of glucosylceramide (37). There is clearly a selective incorporation of sphingolipid into the forming myelin membrane that excludes glucosylceramide and enriches for galactosylceramide and its sulfated derivative, sulfatide. It is notable that deletion of glucosylceramide synthesis in oligodendrocytes does not have obvious effects on myelination (37) so the role of glucosylceramide in oligodendrocyte function remains unresolved.

We carefully examined the regulation of serine palmitoyltransferase (SPT). As the rate-limiting enzyme in sphingolipid biosynthesis it would be expected that SPT activity would increase dramatically during myelination. Our activity, protein and RNA measurements revealed the limitation of analyzing total brain samples in which the number of many cell types are expanding during the period that we analyzed. We found that SPT enzymatic activity measured in whole brain peaks early, day 9 after birth, just at the initiation of myelination rather than continuing to rise as myelin is produced. The activity measurement was consistent with the decrease of SPTLC2 and 3 at the protein and RNA levels during this later period. Elevated SPT activity prior to myelination is likely to reflect the large ganglioside synthesis that accompanies the generation of neurons, axonal elongation and synaptogenesis; as gangliosides are major components of the neuronal membranes (57). The discrepancy between the decline of SPT activity measured in total brain during the period of rising levels of myelin production was resolved by measurement of SPT protein and mRNA in in myelinating oligodendrocytes, a relatively minor cell type in the brain. Measurements of the major SPT subunits at the RNA level in oligodendrocytes revealed that SPTLC1 and −2 levels are elevated at day 16, when myelin-specific sphingolipids are rapidly increasing. Unfortunately there was not sufficient oligodendrocyte material to perform activity measurements to confirm that the SPTLC1 and −2 levels reflect SPT activity.

Critically, activity of SPT with regard to overall activity and substrate and product specificity is dependent on the composition of the complex (reviewed in (58)). This includes whether the active site-containing subunit is SPTLC2 or −3, on presence of the small subunits, ssSPTa or – b, and on the regulatory proteins, the ORMDLs. The predominant acyl-CoA substrate for SPT is the 16 carbon form, palmitoyl-CoA, and this is the preference of the SPTLC1/2/ssSPTa complex, producing d18 sphingoid base-containing sphingolipids, the major forms that we and others have generally measured. Substitution of SPTLC3 for SPTLC2 promotes the use of the 14 carbon acyl-CoA and thus generates d16 sphingoid bases (59). Indeed we find that sphingolipids containing the d16:1 sphingoid base are observed in all sphingolipids that we measured, albeit at low levels (Figure 9, Panel A). The levels of the d16:1-containing sphingolipids generally follow that of the d18:1 sphingolipids and are considerably lower. The significance of the distinct and dramatic peak in SPTLC3 expression at the beginning of the myelination process, observed both in the total brain and oligodendrocyte samples, is not clear. While it may influence the generation of d16 sphingolipids, there may be an as yet unappreciated function of the SPTLC3 subunit that is critical for last stages of oligodendrocyte maturation and the initiation of rapid myelin production. The small subunits, in addition to increasing the overall activity of the SPTLC1/2 dimer also affect the substrate preference of SPT (14). In the context of SPTLC1/2, ssSPTb promotes the use of longer chain fatty acyl-CoA substrates and in the context of SLPTLC1/3, ssSPTb appears to reduce the overall specificity of acyl-CoA utilization. We noted a rise in ssSPTb levels both in the total brain and in oligodendrocyte samples during the peak period of myelination. Correspondingly, the levels of d20:1-containing sphingolipids in oligodendrocytes rise during this period (Figure 9, Panel B). The levels of these lipids are appreciable, up to 8% of the total. Future studies manipulating the levels of ssSPTb will be required to determine the functional consequences of these changes. Indeed, a mouse mutant harboring a hypermorphic mutation in ssSPTb increases levels of d20 sphingolipids and results in a neurodegeneration phenotype (60). Our observation that the composition of the SPT complex changes during oligodendrocyte differentiation and myelination indicates that there are subtle alterations of the sphingolipids that are produced that may have important consequences for myelin assembly and/or function.

The ORMDLs are homeostatic regulators of SPT, inhibiting activity of this enzyme as cellular sphingolipids rise (reviewed in (17)). The ORMDL/SPT complex responds directly to levels of ceramide (61). There are three ORMDL isoforms. In cultured cell lines the three isoforms are functionally redundant (15). This system is thought to maintain ceramide levels at sub-apoptotic levels by ensuring that SPT stimulation of ceramide production does not overwhelm enzymes downstream of ceramide. This would be particularly important in cells, such as the myelinating oligodendrocyte, in which high levels of sphingolipid are being produced. Indeed it has recently been reported that a whole animal double knockout of ORMDLs 1 and 3 results in profound myelin aberration (38). These animals exhibit reduced numbers of myelinated axons and, in the remaining myelin, what appears to be overproduction of the myelin membrane. The authors speculate that unregulated sphingolipid production is both cytotoxic to oligodendrocytes and results in an unregulated myelin production.

We find that in the maturing oligodendrocytes the total level of ORMDL protein is relatively constant. (Figure 8, Panel A). It should be noted that the close homology of the three ORMDL isoforms precludes distinguishing individual isoforms by currently available antibodies. However analysis of mRNA levels indicated that the contribution of the individual isoforms changes as oligodendrocytes mature. Levels of ORMDL1 gradually increase with oligodendrocyte maturation, while ORMDL2 rises sharply in the cells from 8-day-old pups. These cells consist predominantly of pre-oligodendrocytes ready to undergo the last stage of oligodendrocyte maturation into actively myelinating cells. In contrast, levels of ORMDL3 drop somewhat during this period. It remains to be determined how the change in ORMDL isoform composition affects SPT regulation and how these results relate to the observations of Proia and colleagues. One possibility is that each isoform establishes a different ceramide set-point or responds to a different spectrum of ceramide molecular species. It is also possible that the ORMDLs have regulatory functions on other aspects of sphingolipid metabolism.

These studies emphasize that although the general outline of sphingolipid content of sphingolipid in the myelin membrane has been known for decades, little is understood about the biosynthesis and incorporation of these lipids into the forming myelin membrane. Important gaps in our knowledge remain concerning how the levels of individual lipids are produced in the right proportions both in relationship to each other and to the protein components of myelin. The mechanism that transports both the protein and lipid components of myelin from their sites of synthesis into the forming myelin membrane is also poorly understood. An understanding of the details of myelin biosynthesis will be crucial to the development of treatment of demyelinating diseases by either preventing demyelination or stimulating remyelination.

## Abbreviations

SPT: serine palmitoyltransferase;
UGCG: UDP-glucose ceramide glucosyltransferase;
UGT8: UDP-glycosyltransferase 8;
Gal3T: Galactose-3-O-sulfotransferase;
MBP: myelin basic protein;
MOG: myelin oligodendrocyte glycoprotein;
PLP: myelin proteolipid protein;
MAG: myelin associated glycoprotein;
CERT: ceramide transfer protein;
CER: ceramide;
SM: sphingomyelin;
GluCer: glucosylceramide;
GalCer: galactosylceramide.

## Acknowledgements

Supported by NIH R01HL131340, an award from the Presidential Research Quest Fund of Virginia Commonwealth University to BW, and partially by GrantRG 1401-2891 from the National Multiple Sclerosis Society to CS-B. The VCU Lipidomics/Metabolomics Core is supported by the NIH-NCI Cancer Center Support Grant P30 CA016059 to the VCU Massey Cancer Center, as well as a shared resource grant (S10RR031535) from the National Institutes of Health.

The content of this manuscript is solely the responsibility of the authors and does not necessarily represent the official views of the National Institutes of Health.

## Literature Cited

1. Stadelmann, C., Timmler, S., Barrantes-Freer, A., and Simons, M. (2019) Myelin in the Central Nervous System: Structure, Function, and Pathology. Physiological reviews 99, 1381–1431

2. Norton, W. T., and Poduslo, S. E. (1973) Myelination in rat brain: changes in myelin composition during brain maturation. J Neurochem 21, 759–773

3. Camargo, N., Goudriaan, A., van Deijk, A. F., Otte, W. M., Brouwers, J. F., Lodder, H., Gutmann, D. H., Nave, K. A., Dijkhuizen, R. M., Mansvelder, H. D., Chrast, R., Smit, A. B., and Verheijen, M. H. G. (2017) Oligodendroglial myelination requires astrocyte-derived lipids. PLoS biology 15, e1002605

4. Ozgen, H., Baron, W., Hoekstra, D., and Kahya, N. (2016) Oligodendroglial membrane dynamics in relation to myelin biogenesis. Cellular and molecular life sciences : CMLS 73, 3291–3310

5. Bosio, A., Binczek, E., Haupt, W. F., and Stoffel, W. (1998) Composition and biophysical properties of myelin lipid define the neurological defects in galactocerebroside- and sulfatide-deficient mice. J Neurochem 70, 308–315

6. Eckhardt, M. (2008) The role and metabolism of sulfatide in the nervous system. Molecular neurobiology 37, 93–103

7. Marcus, J., Dupree, J. L., and Popko, B. (2002) Myelin-associated glycoprotein and myelin galactolipids stabilize developing axo-glial interactions. J Cell Biol 156, 567–577

8. Marcus, J., Honigbaum, S., Shroff, S., Honke, K., Rosenbluth, J., and Dupree, J. L. (2006) Sulfatide is essential for the maintenance of CNS myelin and axon structure. Glia 53, 372–381

9. Palavicini, J. P., Wang, C., Chen, L., Ahmar, S., Higuera, J. D., Dupree, J. L., and Han, X. (2016) Novel molecular insights into the critical role of sulfatide in myelin maintenance/function. J Neurochem 139, 40–54

10. Bosio, A., Binczek, E., and Stoffel, W. (1996) Functional breakdown of the lipid bilayer of the myelin membrane in central and peripheral nervous system by disrupted galactocerebroside synthesis. Proceedings of the National Academy of Sciences of the United States of America 93, 13280–13285

11. Coetzee, T., Dupree, J. L., and Popko, B. (1998) Demyelination and altered expression of myelin-associated glycoprotein isoforms in the central nervous system of galactolipid-deficient mice. Journal of neuroscience research 54, 613–622

12. Honke, K., Hirahara, Y., Dupree, J., Suzuki, K., Popko, B., Fukushima, K., Fukushima, J., Nagasawa, T., Yoshida, N., Wada, Y., and Taniguchi, N. (2002) Paranodal junction formation and spermatogenesis require sulfoglycolipids. Proceedings of the National Academy of Sciences of the United States of America 99, 4227–4232

13. Harrison, P. J., Dunn, T. M., and Campopiano, D. J. (2018) Sphingolipid biosynthesis in man and microbes. Natural product reports 35, 921–954

14. Han, G., Gupta, S. D., Gable, K., Niranjanakumari, S., Moitra, P., Eichler, F., Brown, R. H., Jr., Harmon, J. M., and Dunn, T. M. (2009) Identification of small subunits of mammalian serine palmitoyltransferase that confer distinct acyl-CoA substrate specificities. Proc. Natl. Acad. Sci. U. S. A 106, 8186–8191

15. Siow, D. L., and Wattenberg, B. W. (2012) Mammalian ORMDL Proteins Mediate the Feedback Response in Ceramide Biosynthesis. J. Biol. Chem 287, 40198–40204

16. van, E. G., Birk, R., Brenner-Weiss, G., Schmidt, R. R., and Sandhoff, K. (1990) Modulation of sphingolipid biosynthesis in primary cultured neurons by long chain bases. J. Biol. Chem 265, 9333–9339

17. Davis, D., Kannan, M., and Wattenberg, B. (2018) Orm/ORMDL proteins: Gate guardians and master regulators. Advances in biological regulation

18. Stiban, J., Tidhar, R., and Futerman, A. H. (2010) Ceramide synthases: roles in cell physiology and signaling. Adv. Exp. Med. Biol 688:60–71., 60–71

19. Chrast, R., Saher, G., Nave, K. A., and Verheijen, M. H. (2011) Lipid metabolism in myelinating glial cells: lessons from human inherited disorders and mouse models. J Lipid Res 52, 419–434

20. Coant, N., Sakamoto, W., Mao, C., and Hannun, Y. A. (2017) Ceramidases, roles in sphingolipid metabolism and in health and disease. Advances in biological regulation 63, 122–131

21. Maceyka, M., Harikumar, K. B., Milstien, S., and Spiegel, S. (2012) Sphingosine-1-phosphate signaling and its role in disease. Trends Cell Biol 22, 50–60

22. Colello, R. J., and Sato-Bigbee, C. (2001) Purification of oligodendrocytes and their progenitors using immunomagnetic separation and Percoll gradient centrifugation. Current protocols in neuroscience Chapter 3, Unit 3.12

23. Lloyd-Evans, E., Pelled, D., Riebeling, C., Bodennec, J., de-Morgan, A., Waller, H., Schiffmann, R., and Futerman, A. H. (2003) Glucosylceramide and glucosylsphingosine modulate calcium mobilization from brain microsomes via different mechanisms. J Biol Chem 278, 23594–23599

24. Aranda, P. S., LaJoie, D. M., and Jorcyk, C. L. (2012) Bleach gel: a simple agarose gel for analyzing RNA quality. Electrophoresis 33, 366–369

25. Schagger, H. (2006) Tricine-SDS-PAGE. Nature protocols 1, 16–22

26. Ren, J., Snider, J., Airola, M. V., Zhong, A., Rana, N. A., Obeid, L. M., and Hannun, Y. A. (2018) Quantification of 3-ketodihydrosphingosine using HPLC-ESI-MS/MS to study SPT activity in yeast Saccharomyces cerevisiae. J Lipid Res 59, 162–170

27. Campagnoni, C. W., Carey, G. D., and Campagnoni, A. T. (1978) Synthesis of myelin basic proteins in the developing mouse brain. Arch Biochem Biophys 190, 118–125

28. Carson, J. H., Nielson, M. L., and Barbarese, E. (1983) Developmental regulation of myelin basic protein expression in mouse brain. Developmental biology 96, 485–492

29. Fewou, S. N., Bussow, H., Schaeren-Wiemers, N., Vanier, M. T., Macklin, W. B., Gieselmann, V., and Eckhardt, M. (2005) Reversal of non-hydroxy:alpha-hydroxy galactosylceramide ratio and unstable myelin in transgenic mice overexpressing UDP-galactose:ceramide galactosyltransferase. J Neurochem 94, 469–481

30. Zoller, I., Meixner, M., Hartmann, D., Bussow, H., Meyer, R., Gieselmann, V., and Eckhardt, M. (2008) Absence of 2-hydroxylated sphingolipids is compatible with normal neural development but causes late-onset axon and myelin sheath degeneration. The Journal of neuroscience : the official journal of the Society for Neuroscience 28, 9741–9754

31. Garbay, B., and Cassagne, C. (1994) Expression of the ceramide galactosyltransferase gene during myelination of the mouse nervous system. Comparison with the genes encoding myelin basic proteins, choline kinase and CTP:phosphocholine cytidylyltransferase. Brain research. Developmental brain research 83, 119–124

32. Koul, O., Chou, K. H., and Jungalwala, F. B. (1980) UDP-galactose-ceramide galactosyltransferase in rat brain myelin subfractions during development. The Biochemical journal 186, 959–969

33. Hannun, Y. A., and Obeid, L. M. (1995) Ceramide: an intracellular signal for apoptosis. Trends in biochemical sciences 20, 73–77

34. Dasgupta, S., and Ray, S. K. (2017) Diverse Biological Functions of Sphingolipids in the CNS: Ceramide and Sphingosine Regulate Myelination in Developing Brain but Stimulate Demyelination during Pathogenesis of Multiple Sclerosis. Journal of neurology and psychology 5

35. Karunakaran, I., and van Echten-Deckert, G. (2017) Sphingosine 1-phosphate - A double edged sword in the brain. Biochim Biophys Acta Biomembr 1859, 1573–1582

36. Semple, B. D., Blomgren, K., Gimlin, K., Ferriero, D. M., and Noble-Haeusslein, L. J. (2013) Brain development in rodents and humans: Identifying benchmarks of maturation and vulnerability to injury across species. Progress in neurobiology 106-107, 1–16

37. Saadat, L., Dupree, J. L., Kilkus, J., Han, X., Traka, M., Proia, R. L., Dawson, G., and Popko, B. (2010) Absence of oligodendroglial glucosylceramide synthesis does not result in CNS myelin abnormalities or alter the dysmyelinating phenotype of CGT-deficient mice. Glia 58, 391–398

38. Clarke, B. A., Majumder, S., Zhu, H., Lee, Y. T., Kono, M., Li, C., Khanna, C., Blain, H., Schwartz, R., Huso, V. L., Byrnes, C., Tuymetova, G., Dunn, T. M., Allende, M. L., and Proia, R. L. (2019) The Ormdl genes regulate the sphingolipid synthesis pathway to ensure proper myelination and neurologic function in mice. eLife 8, e51067

39. Zoller, I., Bussow, H., Gieselmann, V., and Eckhardt, M. (2005) Oligodendrocyte-specific ceramide galactosyltransferase (CGT) expression phenotypically rescues CGT-deficient mice and demonstrates that CGT activity does not limit brain galactosylceramide level. Glia 52, 190–198

40. Jimenez-Rojo, N., Garcia-Arribas, A. B., Sot, J., Alonso, A., and Goni, F. M. (2014) Lipid bilayers containing sphingomyelins and ceramides of varying N-acyl lengths: a glimpse into sphingolipid complexity. Biochimica et biophysica acta 1838, 456–464

41. Kolesnick, R. N., Goni, F. M., and Alonso, A. (2000) Compartmentalization of ceramide signaling: physical foundations and biological effects. Journal of cellular physiology 184, 285–300

42. Ben-David, O., Pewzner-Jung, Y., Brenner, O., Laviad, E. L., Kogot-Levin, A., Weissberg, I., Biton, I. E., Pienik, R., Wang, E., Kelly, S., Alroy, J., Raas-Rothschild, A., Friedman, A., Brugger, B., Merrill, A. H., Jr., and Futerman, A. H. (2011) Encephalopathy caused by ablation of very long acyl chain ceramide synthesis may be largely due to reduced galactosylceramide levels. J Biol Chem 286, 30022–30033

43. Schmitt, S., Castelvetri, L. C., and Simons, M. (2015) Metabolism and functions of lipids in myelin. Biochimica et biophysica acta 1851, 999–1005

44. Novgorodov, S. A., Chudakova, D. A., Wheeler, B. W., Bielawski, J., Kindy, M. S., Obeid, L. M., and Gudz, T. I. (2011) Developmentally regulated ceramide synthase 6 increases mitochondrial Ca2+ loading capacity and promotes apoptosis. J. Biol. Chem 286, 4644–4658

45. Hannun, Y. A., and Obeid, L. M. (2002) The Ceramide-centric Universe of Lipid-mediated Cell Regulation: Stress Encounters of the Lipid Kind. Journal of Biological Chemistry 277, 25847–25850

46. Hirahara, Y., Wakabayashi, T., Mori, T., Koike, T., Yao, I., Tsuda, M., Honke, K., Gotoh, H., Ono, K., and Yamada, H. (2017) Sulfatide species with various fatty acid chains in oligodendrocytes at different developmental stages determined by imaging mass spectrometry. J Neurochem 140, 435–450

47. Becker, I., Wang-Eckhardt, L., Yaghootfam, A., Gieselmann, V., and Eckhardt, M. (2008) Differential expression of (dihydro)ceramide synthases in mouse brain: oligodendrocyte-specific expression of CerS2/Lass2. Histochemistry and cell biology 129, 233–241

48. Imgrund, S., Hartmann, D., Farwanah, H., Eckhardt, M., Sandhoff, R., Degen, J., Gieselmann, V., Sandhoff, K., and Willecke, K. (2009) Adult ceramide synthase 2 (CERS2)-deficient mice exhibit myelin sheath defects, cerebellar degeneration, and hepatocarcinomas. J Biol Chem 284, 33549–33560

49. Mesicek, J., Lee, H., Feldman, T., Jiang, X., Skobeleva, A., Berdyshev, E. V., Haimovitz-Friedman, A., Fuks, Z., and Kolesnick, R. (2010) Ceramide synthases 2, 5, and 6 confer distinct roles in radiation-induced apoptosis in HeLa cells. Cell Signal 22, 1300–1307

50. Sassa, T., Suto, S., Okayasu, Y., and Kihara, A. (2012) A shift in sphingolipid composition from C24 to C16 increases susceptibility to apoptosis in HeLa cells. Biochimica et biophysica acta 1821, 1031–1037

51. Senkal, C. E., Ponnusamy, S., Manevich, Y., Meyers-Needham, M., Saddoughi, S. A., Mukhopadyay, A., Dent, P., Bielawski, J., and Ogretmen, B. (2011) Alteration of ceramide synthase 6/C16-ceramide induces activating transcription factor 6-mediated endoplasmic reticulum (ER) stress and apoptosis via perturbation of cellular Ca2+ and ER/Golgi membrane network. J Biol Chem 286, 42446–42458

52. Wattenberg, B. W. (2018) The long and the short of ceramides. J Biol Chem 293, 9922–9923

53. Yamaji, T., and Hanada, K. (2015) Sphingolipid metabolism and interorganellar transport: localization of sphingolipid enzymes and lipid transfer proteins. Traffic 16, 101–122

54. Kudo, N., Kumagai, K., Tomishige, N., Yamaji, T., Wakatsuki, S., Nishijima, M., Hanada, K., and Kato, R. (2008) Structural basis for specific lipid recognition by CERT responsible for nonvesicular trafficking of ceramide. Proceedings of the National Academy of Sciences of the United States of America 105, 488–493

55. Kumagai, K., Yasuda, S., Okemoto, K., Nishijima, M., Kobayashi, S., and Hanada, K. (2005) CERT mediates intermembrane transfer of various molecular species of ceramides. J Biol Chem 280, 6488–6495

56. Funato, K., Riezman, H., and Muniz, M. (2019) Vesicular and non-vesicular lipid export from the ER to the secretory pathway. Biochimica et biophysica acta. Molecular and cell biology of lipids 1865, s1388–1981

57. Yu, R. K., Nakatani, Y., and Yanagisawa, M. (2009) The role of glycosphingolipid metabolism in the developing brain. J Lipid Res 50 Suppl, S440–445

58. Lowther, J., Naismith, J. H., Dunn, T. M., and Campopiano, D. J. (2012) Structural, mechanistic and regulatory studies of serine palmitoyltransferase. Biochem. Soc. Trans 40, 547–554

59. Hornemann, T., Penno, A., Rütti, M. F., Ernst, D., Kivrak-Pfiffner, F., Rohrer, L., and von Eckardstein, A. (2009) The SPTLC3 subunit of serine palmitoyltransferase generates short chain sphingoid bases. The Journal of biological chemistry 284, 26322–26330

60. Zhao, L., Spassieva, S., Gable, K., Gupta, S. D., Shi, L.-Y., Wang, J., Bielawski, J., Hicks, W. L., Krebs, M. P., Naggert, J., Hannun, Y. A., Dunn, T. M., and Nishina, P. M. (2015) Elevation of 20-carbon long chain bases due to a mutation in serine palmitoyltransferase small subunit b results in neurodegeneration. Proceedings of the National Academy of Sciences of the United States of America 112, 12962–12967

61. Davis, D. L., Gable, K., Suemitsu, J., Dunn, T. M., and Wattenberg, B. W. (2019) The ORMDL/Orm-serine palmitoyltransferase (SPT) complex is directly regulated by ceramide: Reconstitution of SPT regulation in isolated membranes. J Biol Chem 294, 5146–5156

